# Population replacement gene drive characteristics for malaria elimination in a range of seasonal transmission settings: a modeling study

**DOI:** 10.1101/2021.11.01.466856

**Authors:** Shirley Leung, Nikolai Windblicher, Edward Wenger, Caitlin Bever, Prashanth Selvaraj

## Abstract

Genetically engineering mosquitoes is a promising new vector control strategy to reinvigorate the fight against malaria in Sub-Saharan Africa. Using an agent-based model of malaria transmission with vector genetics, we examine the impacts of releasing population-replacement gene drive mosquitoes on malaria transmission and quantify the gene drive system parameters required to achieve local elimination within a spatially-resolved, seasonal Sahelian setting. We evaluate the performance of two different gene drive systems: “classic” and “integral.” Various transmission regimes (low, moderate, and high - corresponding to annual entomological inoculation rates of 10, 30, and 80 infectious bites per person) and other simultaneous interventions, including deployment of insecticide-treated nets (ITNs) and passive healthcare seeking, are also simulated. Local elimination probabilities decreased with pre-existing population target site resistance frequency, increased with transmission-blocking effectiveness of the introduced antiparasitic gene and drive efficiency, and were context dependent with respect to fitness costs associated with the introduced gene. Of the four parameters, transmission-blocking effectiveness may be the most important to focus on for improvements to future gene drive strains because a single release of classic gene drive mosquitoes is likely to locally eliminate malaria in low to moderate transmission settings only when transmission-blocking effectiveness is very high (above ^~^80-90%). However, simultaneously deploying ITNs and releasing integral rather than classic gene drive mosquitoes significantly boosts elimination probabilities, such that elimination remains highly likely in low to moderate transmission regimes down to transmission-blocking effectiveness values as low as ^~^50% and in high transmission regimes with transmission-blocking effectiveness values above ^~^80-90%. Thus, a single release of currently achievable population replacement gene drive mosquitoes, in combination with traditional forms of vector control, can likely locally eliminate malaria in low to moderate transmission regimes within the Sahel. In a high transmission regime, higher levels of transmission-blocking effectiveness than are currently available may be required.

**Author summary:** Malaria remains a significant health burden in Sub-Saharan Africa. The mass deployment of insecticide-treated nets and antimalarial drugs have drastically reduced malaria incidence, but insecticide and drug resistance threaten to stall these efforts. The genetic engineering of mosquito populations is a promising new vector control strategy to reinvigorate the fight against malaria. Releases of engineered gene drive mosquitoes that can spread introduced antimalarial genes quickly throughout the mosquito population may be a particularly effective new method for reducing malaria transmission. Important questions arise, however, about how well these gene drive systems must work in order to deliver substantial reductions in transmission. Here we use a spatial model of individual humans and vectors to simulate the effects of releasing gene drive mosquitoes with antimalarial properties on malaria transmission in a Sahelian setting. We quantify the gene drive system parameters required to achieve local elimination and find that when deployed in combination with traditional forms of vector control, a single release of gene drive mosquitoes with realistically achievable characteristics is highly likely to locally eliminate malaria in low to moderate transmission regimes. In a high transmission regime, improved strains of gene drive mosquitoes may be required. In all settings, releasing gene drive mosquitoes with antimalarial properties helps create a window of opportunity during which malaria prevalence is suppressed and other interventions can be ramped up to achieve elimination, even when a single gene drive mosquito release by itself cannot.

## Introduction

Malaria remains a significant health burden in Sub-Saharan Africa (SSA) despite many decades of effort to eliminate the disease [1]. More recently, since the early 2000s, the scale up and mass deployment of long-lasting insecticide-treated nets, indoor residual spraying of insecticides, and antimalarial drugs have drastically reduced malaria incidence [2]. However, this existing set of tools is unlikely to bring about eradication [[3]]. Drug and insecticide resistance further threaten to stall these malaria control efforts [4–7]. New strategies and technologies will therefore be needed to achieve elimination in SSA. The genetic engineering of mosquito populations is a promising new vector control strategy to reinvigorate the fight against malaria and potentially lead to elimination.

Indeed, releases of genetically modified (GM) sterile male mosquitoes have been used to successfully suppress *Aedes aegypti* vector populations [8–12]. This method is expensive, however, and requires frequent, large-scale releases, known as inundation. Releases of GM gene drive mosquitoes, in contrast, are predicted to be a cost-effective and longer-lasting alternative requiring far fewer and smaller releases [13,14]. Mosquitoes engineered with gene drive systems can copy specified genes from one chromosome to another in germline cells, ensuring that these genes are passed onto their offspring at higher than Mendelian inheritance rates and therefore rapidly spread through a population even if there are associated fitness costs [15].

Gene drive mosquito releases can either aim to reduce (population suppression) or to modify (population replacement) a given vector population [15]. Population replacement gene drive systems are the focus of this study and they consist of a driver gene that enables the copying of both itself and an effector gene, which in turn confers desired phenotypic traits. The driver gene encodes a guide RNA and an endonuclease, such as Cas9, that together recognize and cut specified DNA sequences present in the wildtype mosquito population. Within mosquitoes that are heterozygous for the wildtype and drive or effector alleles, the cut wildtype chromosome uses its intact drive or effector-containing sister chromosome as a template for repairing itself, copying over the intact chromosome’s drive or effector-containing DNA in the process through homology-directed repair (HDR) [16].

Population replacement may be desirable in locations where the ecological effects of removing a mosquito species are not well known. For population replacement applied to malaria reduction or elimination, many potential effector genes have been shown to impair development of *Plasmodium* parasites by *Anopheles* mosquitoes. These include genes that code for immune system activators, peptides that neutralize *Plasmodium* parasites in the mosquito midgut or salivary glands, and others [17–23].

A number of important questions about population replacement drives require further investigation. How effective do these effector genes have to be in order to deliver substantial reductions in malaria transmission? Can elimination be achieved even with imperfect transmission blocking traits? If there are significant fitness costs associated with expressing the effector, can it nonetheless propagate quickly within the vector population? Questions also arise around the required efficiency of the driver gene and gene drive system itself to achieve elimination. For example, the process of copying the effector gene from one chromosome to another is not always successful. After cutting, DNA can sometimes undergo alternative repair pathways that do not result in accurate copying of the drive or effector-containing DNA on the sister chromosome. Non-homologous end-joining (NHEJ), microhomology mediated end-joining, or incomplete HDR may occur instead with different probabilities, generating “resistant” alleles that do not contain the desired drive or effector gene but are no longer recognized by the driver endonuclease [24–26]. These resistant alleles may also be present in the wild mosquito population even before introduction of new drive or effector genes [27]. The extent to which the generation and pre-existing presence of these resistant alleles affects the ability of introduced gene drive mosquitoes to eliminate malaria must be better quantified.

Because the potential harms and possible ecological risks associated with releasing gene drive mosquitoes into the wild have not yet been well established, it is not currently feasible or ethical to test such releases in the field. Community understanding, support, and buy-in are also needed before gene drive mosquito releases can proceed [28–30]. Modeling is therefore a key step needed to quantify both the potential benefits and harmful impacts of gene drive mosquito release. Modeling can also help inform the minimum efficacy and genetic parameters required of engineered mosquitoes to achieve substantial public health impacts, thus driving efficient and targeted development of genetically engineered mosquitoes in the laboratory [31].

Here we examine the impacts of releasing malaria transmission-blocking, gene drive mosquitoes in a rural Sahelian setting and quantify the gene drive system characteristics required to achieve elimination. We use an individual-based model of malaria transmission that also resolves agent-based vector genetics and allows for many-to-many mappings of vector genotypes to phenotypes [32]. We quantify the difference in malaria outcomes across a range of transmission settings between releasing two different population replacement gene drive mosquitoes (classic and integral), as well as with and without other forms of vector control. Previous modeling work has focused on understanding changes in vector populations with release of GM mosquitoes without considering other types of vector control and without also examining the downstream effects on malaria transmission within corresponding human populations [31,33–37]. An advantage of our model [32,38] is that it can simulate the effects of gene drive-induced vector population changes on malaria transmission within a realistic human population directly. Because of this added ability, we are able to quantify the gene drive system and other logistical release parameters needed to reach full malaria elimination.

## Methods

### Model overview

Simulations were carried out using EMOD v2.20 [39], which is a mechanistic, agent-based model of *Plasmodium falciparum* malaria transmission that can individually track each mosquito’s movement and feeding pattern as well as each human’s movement, infection, and immune dynamics. Mosquitoes within EMOD go through four life cycle stages: eggs, larvae, immature adults that do not seek hosts or reproduce, and mature adults that do seek hosts and reproduce [40]. While adult female mosquitoes can complete their feeding cycle and lay eggs, the number of eggs that progress to the larval stage is determined by the amount of larval habitat available at a given time, which in turn governs the number of adult vectors that eventually emerge.

Mosquitoes within EMOD contain simulated genomes represented by up to 10 different loci or genes, with up to 8 different alleles per gene. Various phenotypic traits can be assigned to different genotypes, including changes in fecundity, malaria transmissibility, mortality, and insecticide resistance.

When an adult male and female mosquito mate in the model, they each contribute half of the genes belonging to their offspring. During gametogenesis before meiosis is complete, the germline cells within each parent mosquito undergo all necessary gene drive-related changes to their genomes. After completion of all drive-related changes, each parent’s germline cells undergo meiosis and gametes are distributed to offspring according to Mendelian inheritance. Further details regarding the implementation of vector genetics within EMOD are explained in See Selvaraj et al. (2020) [32].

Human agents within EMOD each have their own microsolver to track within-host parasite dynamics and the associated parasitological and clinical immunity that arise from innate and adaptive responses to specific antigens. Parameters associated with this microsolver have been calibrated to reflect transmission in a range of scenarios in Sub-Saharan Africa under different transmission intensities and with or without interventions [41].

#### Modeled region

To capture conditions representative of the Sahel region of SSA, simulations were conducted over a 300 square kilometer region of rural Burkina Faso (Figure 1A). This 300 square kilometer region was divided into 1 km-by-1 km grid cells, each with its own simulated human and vector population. Human population data from the region was obtained from the High Resolution Settlement Layer generated by the Facebook Connectivity Lab and Columbia University’s Center for International Earth Science Information Network [42]. Only grid cells with more than 5 people were included in the simulations, resulting in ^~^3,700 individuals simulated across 150 populated grid cells.

**Figure 1.**
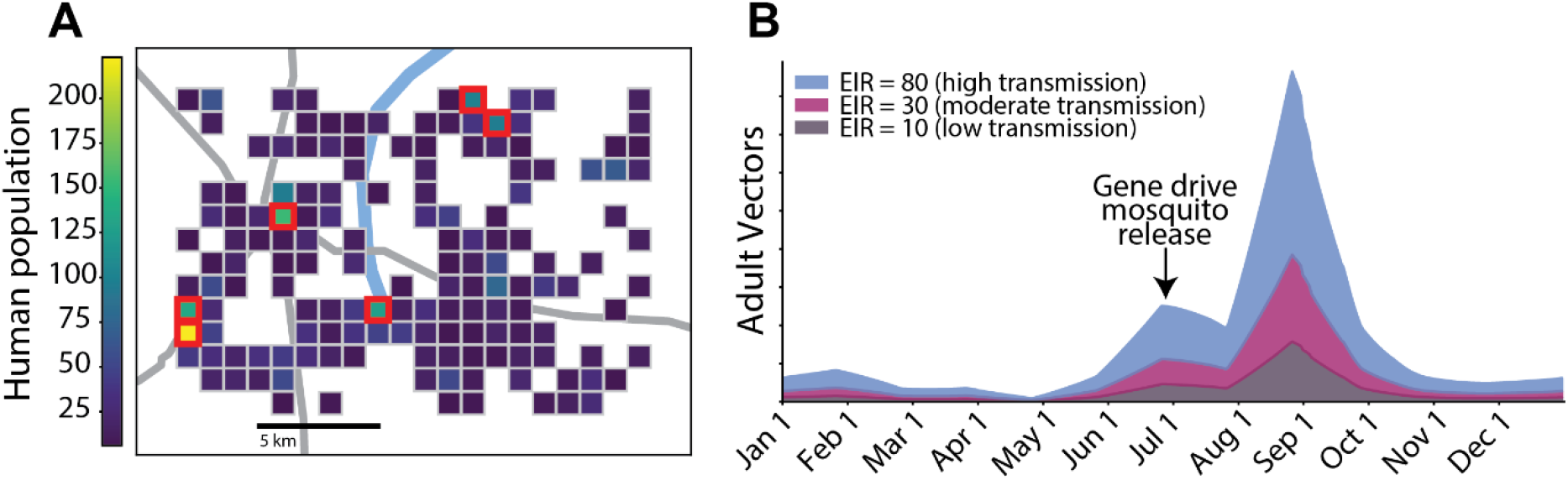
Simulated spatial region and seasonality. (A) Spatial region and grid composed of 150 1 km-by-1 km nodes used for all simulations. Colors denote the human population within each node. In all simulation, 100 male gene drive mosquitoes were released in each of the six most populous nodes (outlined in red), which account for ^~^23% of the human population in the region. (B) Baseline seasonal cycle of adult vector populations within the simulated area before gene drive releases in the three low (annual EIR = 10 infectious bites per person), moderate (annual EIR = 30 infectious bites per person), and high (annual EIR = 80 infectious bites per person) Sahelian transmission regimes simulated here. Gene drive mosquitoes were released on July 1 of the first simulation year in all simulations. ITNS were also deployed on July 1 of the first, fourth, and seventh simulation years in simulations with ITNs.

Vector carrying capacity and initial populations were scaled to human population within each node to ensure that humans have the same probability of being bitten across all grid cells. Only one vector species, *Anopheles gambiae*, was assumed to be present and responsible for all malaria transmission. Characteristic Sahelian seasonality in vector populations was captured by appropriately varying the amount of available larval habitat space over the year (Figure 1B) [41,43–45]. The same seasonal profile of larval habitat space was used in all grid cells and all scenarios. The amplitude of the larval habitat, and in turn mosquito density and biting, was varied to simulate different transmission intensities with annual entomological inoculation rates (EIR) varying between 10 infectious bites per person (reflecting a low transmission setting) to 80 infectious bites per person (reflecting a high transmission setting).

Human migration is simulated by assigning each individual person a daily probability of taking overnight trips to other grid cells. This probability is governed by a gravity model dependent on population in and distance between nodes [46]. The gravity model is calibrated to movements observed in geotagged campaign data [47] and results in an average of 5 overnight trips per person per year. Similar to human migration, vector migration is simulated by assigning each individual mosquito a daily probability of migrating to another grid cell. This probability is governed by a negative exponential distance decay function [48] (Supp. Figure 1). Neither humans nor vectors migrate into or out of the simulated region. There is therefore no importation of malaria from outside of the modelled area.

All scenarios were simulated for 8 years and 20 stochastic realizations were run for each scenario.

#### Modeled interventions

All simulations included treatment with artemether-lumefantrine (AL) for symptomatic cases. Those with severe malaria cases sought treatment 80% of the time within 2 days of symptom onset. Those with clinical, but not severe, cases sought treatment 50% of the time within 3 days of symptom onset. Health-seeking rates are assumed to be the same for all ages.

Some simulations included ITN deployments. Per WHO guidelines [49], ITNs were distributed (Figure 1A) every 3 years at the beginning of the peak season on July 1, covering a random 70% of the population per distribution. To reflect the effects of insecticide resistance, each ITN is set to have a reduced initial vector blocking efficacy of only 60% and killing rate of only 70%. Both blocking and killing rates decay exponentially over time with a decay constant of 2 years and 4 years, respectively. Simulations with ITN deployment alone (that is, those without a gene drive mosquito release) result in elimination probabilities of zero for all transmission regimes (low, moderate, and high) tested here (results not shown).

#### Gene drives

The basic setup of a gene drive system for population replacement involves coupling a driver gene with an anti-malaria effector gene that prevents the mosquito from transmitting malaria. There are, however, multiple ways in which this can be implemented. In what we term “classic” gene drive systems, the driver and effector genes are propagated as a single complex construct and inserted at an arbitrary target site within the genome (Figure 2A). In a more recently conceived “integral” gene drive system, the driver and one or multiple effector genes are separated into distinct molecularly simpler constructs and are then inserted into essential genes [37] (Figure 2B). In this case, the endonuclease produced from the driver mediates homing both of its own gene and the effector gene. A previous compartmental vector model suggests that an integral gene drive system of this type can provide longer-lasting protection from malaria within a vector population than a classical system by both slowing down the generation of resistance alleles and allowing for the Mendelian inheritance of the effector gene even when the driver gene is lost [37]. Here we use EMOD to simulate the release and spread of both classic and integral gene drive mosquitoes.

**Figure 2.**
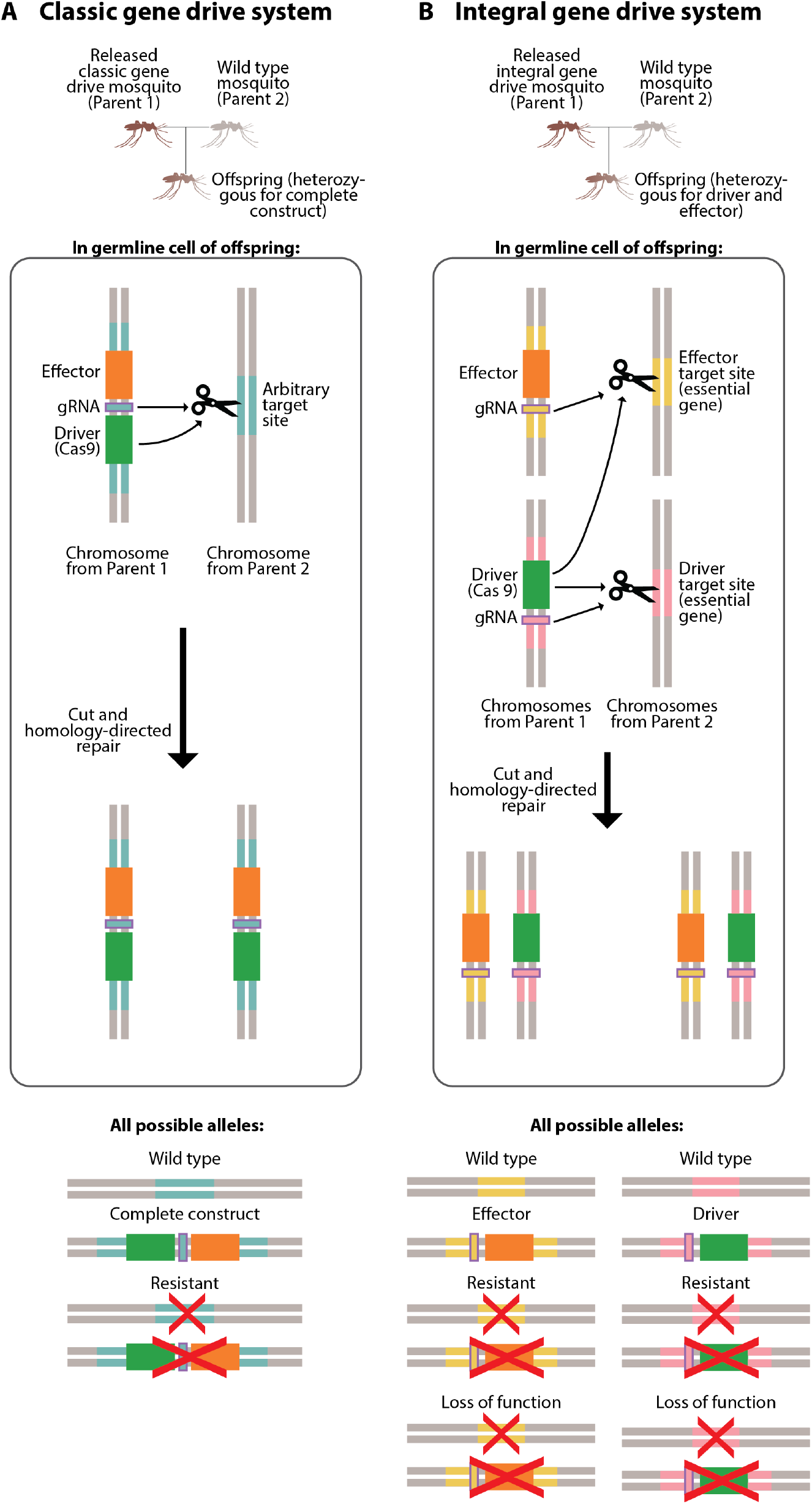
Classic and integral gene drive systems. (A) Classic gene drive system and possible alleles. (B) Integral gene drive system and possible alleles.

In all simulations discussed below, we released 100 male gene drive mosquitoes in each of the 6 most populous nodes (1 km-by-1 km grid cells), for a total of 600 released mosquitoes, on July 1 of the first simulated year. These 6 most populous nodes account for ^~^23% of the total human population in the simulated region. In our classic gene drive release simulations, we allow for the possibility of 3 allele types at the target site locus: wild type, complete construct (drive and effector), and resistant (Figure 2A). Only expression of the complete construct confers anti-pathogenicity and increases fitness cost via enhanced mortality, while only the wild type allele can be recognized and cut by the driver. Resistant alleles may occur naturally in the initial vector population and/or may arise during errors in the homologous copying process; they do not carry any anti-pathogenicity or fitness cost and cannot be recognized by the driver. In our integral gene drive release simulations, we allow for the possibility of 4 allele types at both the driver and effector target site loci: wild type, nuclease (in the case of the driver target site) or effector (in the case of the effector target site), resistant, and loss-of-function (Figure 2B). As with the classic gene drive system, only expression of the nuclease or effector affects vector fitness, while only the wild type alleles can be recognized and cut by the driver. Only expression of the effector confers anti-parasitic properties to the vector. Specific to integral gene drive systems, driver target sites are located within essential, recessive lethal genes; loss-of-function alleles that lead to non-viability in homozygosity can therefore crop up when mutations arise during HDR. Because of conferred non-viability, loss-of-function alleles are disproportionately lost from the population, which consequently increases the proportion of intact, successfully-copied nuclease or effector alleles relative to the classic setup. In all simulations, we assume negligible rates of random mutations at all target sites. Tables 1 and 2 summarize other important classic and integral gene drive system parameters, respectively.

**Table 1.** Classic gene drive system parameters. All genetic parameters used in classic gene drive mosquito release simulations. Default values are representative of and consistent with other published works [31,37,54]. Default values for tested parameters (*d*, *sne*, *rc*, *rr0*) are used on accompanying website visualizations.

| Parameter | Description | Default value | Range |
| --- | --- | --- | --- |
| *d* | Drive efficiency and transmission rate | 1 | 0.9 - 1 |
| *u* | Probability of resistance arising if drive transmission fails | 0.5 | - |
| *lne* | Probability of complete construct loss during homing | 3E-4 | - |
| *sd* | Fitness cost of target site disruption | 0 | - |
| *sne* | Fitness cost of complete construct expression | 0 | 0 - 0.5 |
| *hd* | Dominance coefficient for target site disruption | 0.5 | - |
| *hne* | Dominance coefficient for complete construct expression | 0.5 | - |
| *hrc* | Dominance coefficient for parasite refractoriness | 1 | - |
| *rc* | Homozygous degree of parasite refractoriness | 1 | 0.5 - 1 |
| *rr0* | Initial population target site resistance frequency | 0 | 0 - 0.1 |

**Table 2.**
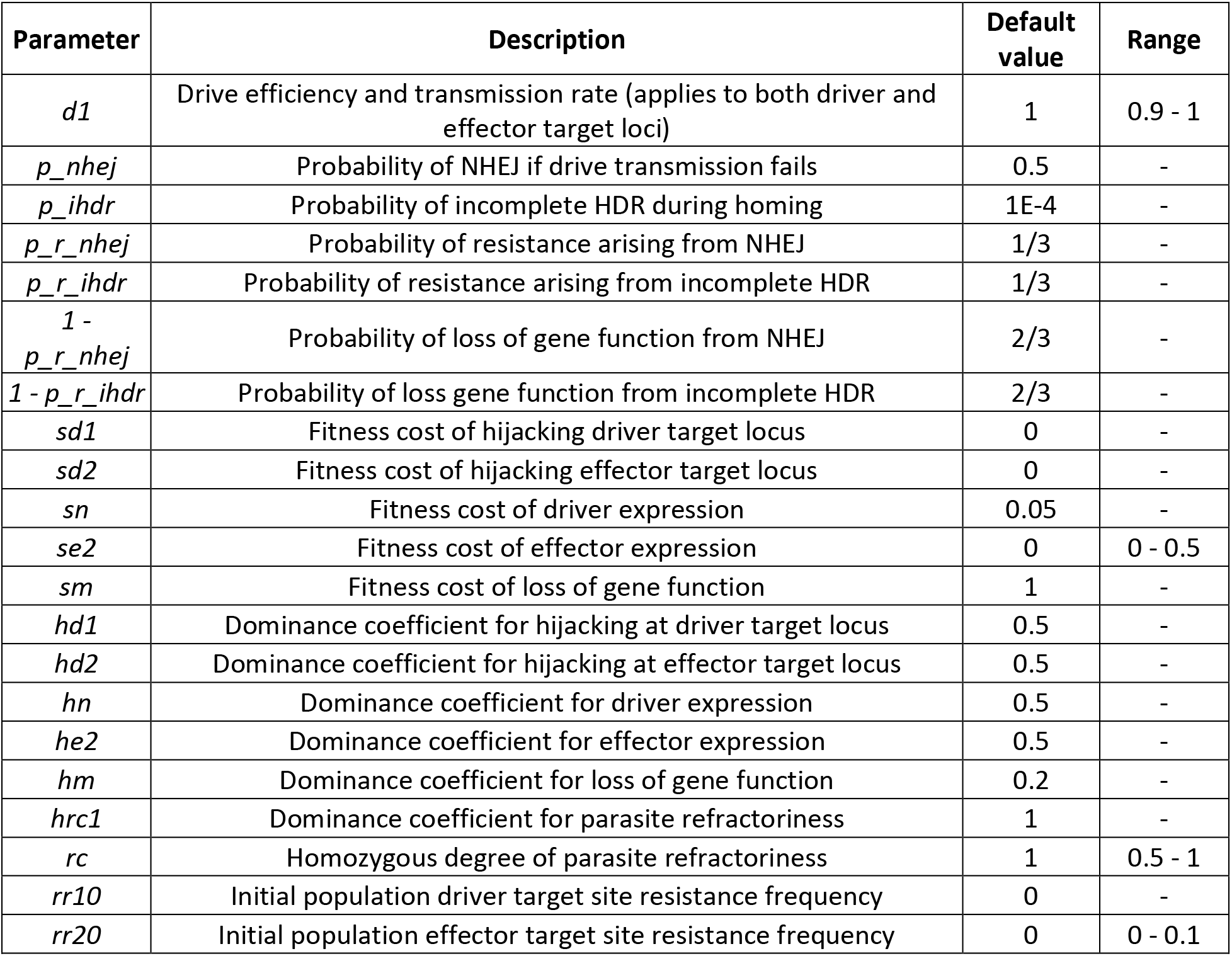
Integral gene drive system parameters. All genetic parameters used in integral gene drive mosquito release simulations. Default values are representative of and consistent with other published works [31,37,54]. Default values for tested parameters (*d1, se2, rc, rr20*) are used on accompanying website visualizations.

| Parameter | Description | Default value | Range |
| --- | --- | --- | --- |
| *d1* | Drive efficiency and transmission rate (applies to both driver and effector target loci) | 1 | 0.9 - 1 |
| *p_nhej* | Probability of NHEJ if drive transmission fails | 0.5 | - |
| *p_ihdr* | Probability of incomplete HDR during homing | 1E-4 | - |
| *p_r_nhej* | Probability of resistance arising from NHEJ | 1/3 | - |
| *p_r_ihdr* | Probability of resistance arising from incomplete HDR | 1/3 | - |
| $1 - p_r_nhej$ | Probability of loss of gene function from NHEJ | 2/3 | - |
| $1 - p_r_ihdr$ | Probability of loss gene function from incomplete HDR | 2/3 | - |
| *sd1* | Fitness cost of hijacking driver target locus | 0 | - |
| *sd2* | Fitness cost of hijacking effector target locus | 0 | - |
| *sn* | Fitness cost of driver expression | 0.05 | - |
| *se2* | Fitness cost of effector expression | 0 | 0 - 0.5 |
| *sm* | Fitness cost of loss of gene function | 1 | - |
| *hd1* | Dominance coefficient for hijacking at driver target locus | 0.5 | - |
| *hd2* | Dominance coefficient for hijacking at effector target locus | 0.5 | - |
| *hn* | Dominance coefficient for driver expression | 0.5 | - |
| *he2* | Dominance coefficient for effector expression | 0.5 | - |
| *hm* | Dominance coefficient for loss of gene function | 0.2 | - |
| *hrc1* | Dominance coefficient for parasite refractoriness | 1 | - |
| *rc* | Homozygous degree of parasite refractoriness | 1 | 0.5 - 1 |
| *rr10* | Initial population driver target site resistance frequency | 0 | - |
| *rr20* | Initial population effector target site resistance frequency | 0 | 0 - 0.1 |

In simulations of both classic and integral gene drive releases, we examine the effects of the following parameters on likelihood of local malaria elimination (defined as malaria prevalence reaching and staying at zero by the end of simulation year 7 within all spatial nodes): the probability of copying over the driver and/or effector genes in the presence of the driver gene (also known as the efficiency of the drive, *d*); the ability of the effector gene to prevent onward malaria transmission in mosquitoes (also known as the transmission-blocking effectiveness of the effector, which is equivalent in either heterozygosity or homozygosity, *rc*); the pre-existing frequency of target site resistance alleles in the population (*rr0* in the classic case; *rr20* at the effector target site in the integral case); and the fitness cost associated with expressing the introduced driver and effector genes, represented by an increase in vector mortality (*sne* in the classic case; *se2* associated with the effector in the integral case).

Because of the high dimensionality of the results, we also created a website with interactive visualizations of simulation output to accompany the figures in this text (Supp. Figure 2), located here: https://gene-drive.bmgf.io. Website users can interactively visualize the effects of all tested parameters on elimination probabilities along with elimination timing, prevalence, vector populations, and allele frequencies over all simulated combinations of gene drive release types, ITN deployments, and transmission regimes. Though we focus primarily on understanding the effects of tested parameters on local elimination probabilities in this text, we highly encourage website users to explore the effects of tested parameters on additional malaria-related variables plotted on the website as well, particularly reductions in prevalence even if elimination is not achieved.

## Results

### Elimination probability decreases with pre-existing resistance, increases with transmission-blocking and drive efficiency, and is context dependent with respect to fitness costs

When conducting a single release of classic gene drive mosquitoes, elimination probabilities increase when transmission-blocking effectiveness (*rc*) increases, drive efficiency (*d*) increases, and pre-existing population target site resistance (*rr0*) decreases over all tested parameter values (Figure 3). Holding all other parameters constant, as transmission-blocking effectiveness increases, each individual mosquito carrying the complete construct in the population is less likely to become infected by the malaria parasite and pass it on to their human hosts. Thus, the higher the transmission-blocking effectiveness, the lower the frequency of vectors that are infectious among the total vector population (Figure 4) and the greater the chance of eliminating malaria within the local population. With all other parameters held constant, an increase in drive efficiency leads to both an earlier and higher peak effector frequency, as the introduced complete construct spreads at super-Mendelian rates through the vector population (Figure 5). Earlier and higher peak effector frequencies at higher drive efficiencies reduce the fraction of the mosquito population that can be infected by malaria parasites, thus increasing local malaria elimination probabilities. The opposite occurs as pre-existing population target site resistance increases with all other parameters held constant. Because target site resistance prevents the spread of the introduced construct, peak effector frequency is reduced and more mosquitoes are able to transmit malaria parasites at higher pre-existing target site resistances (Figure 6). As a result, an increase in pre-existing target site resistance within a vector population reduces the chances of locally eliminating malaria with a single gene drive release. To visualize elimination probabilities with the tested parameters on different axes than those displayed in Figure 3, see https://gene-drive.bmgf.io.

**Figure 3.**
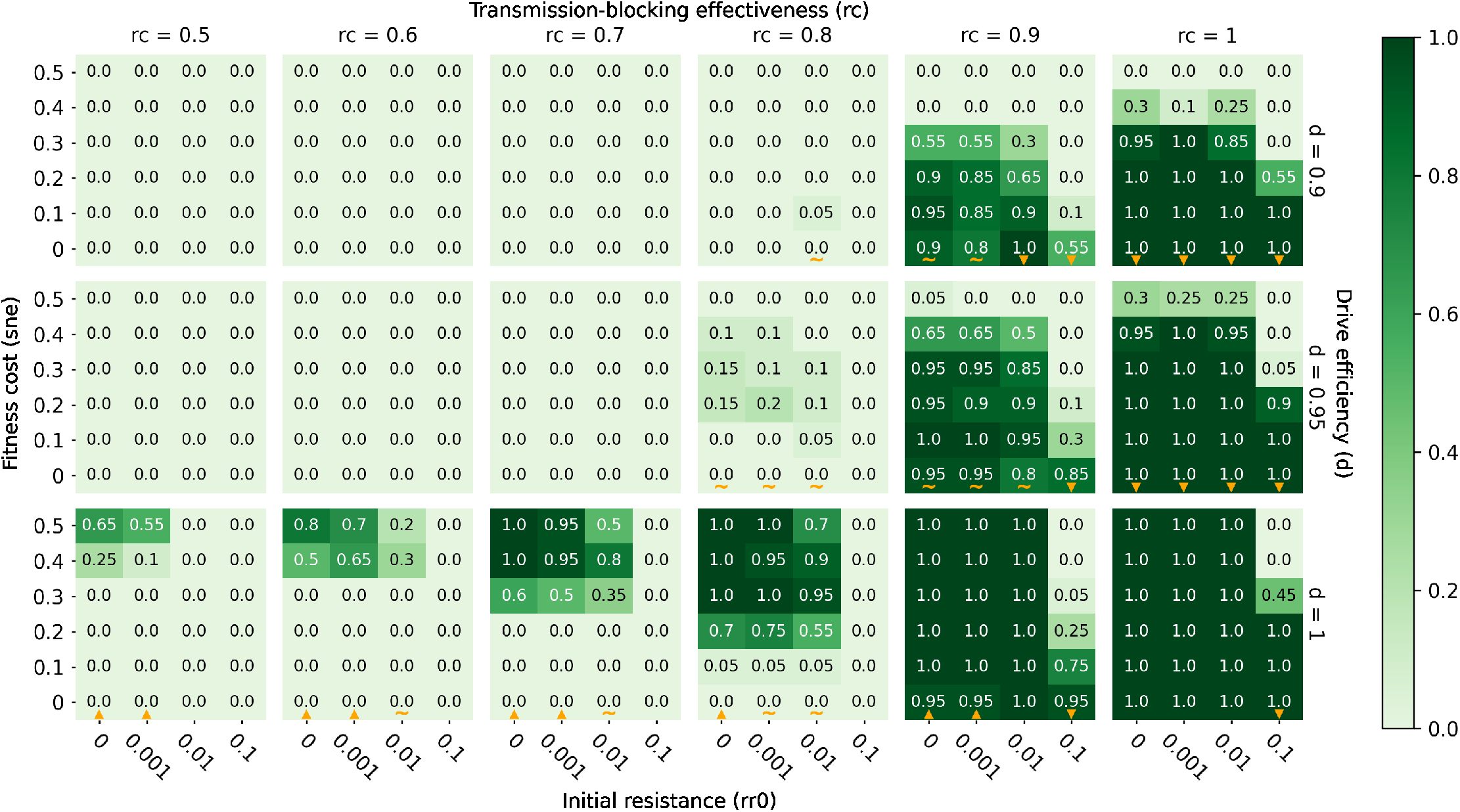
Elimination probabilities after a single release of classic gene drive mosquitoes only in a moderate transmission (annual EIR = 30) regime. Elimination probabilities (computed as the fraction of 20 model realizations in which malaria prevalence reaches and remains at zero by the end of simulation year 7) over a range of transmission-blocking effectiveness (*rc*), drive efficiency (*d*), pre-existing population target site resistance frequency (*rr0*), and mortality-enhancing effector expression fitness cost (*sne*) values. Orange upward and downward-pointing orange triangles denote columns along which elimination probabilities increase and decrease with increasing fitness cost, respectively. Orange tildes denote columns along which elimination probabilities first increase and then decrease with increasing fitness cost.

**Figure 4.**
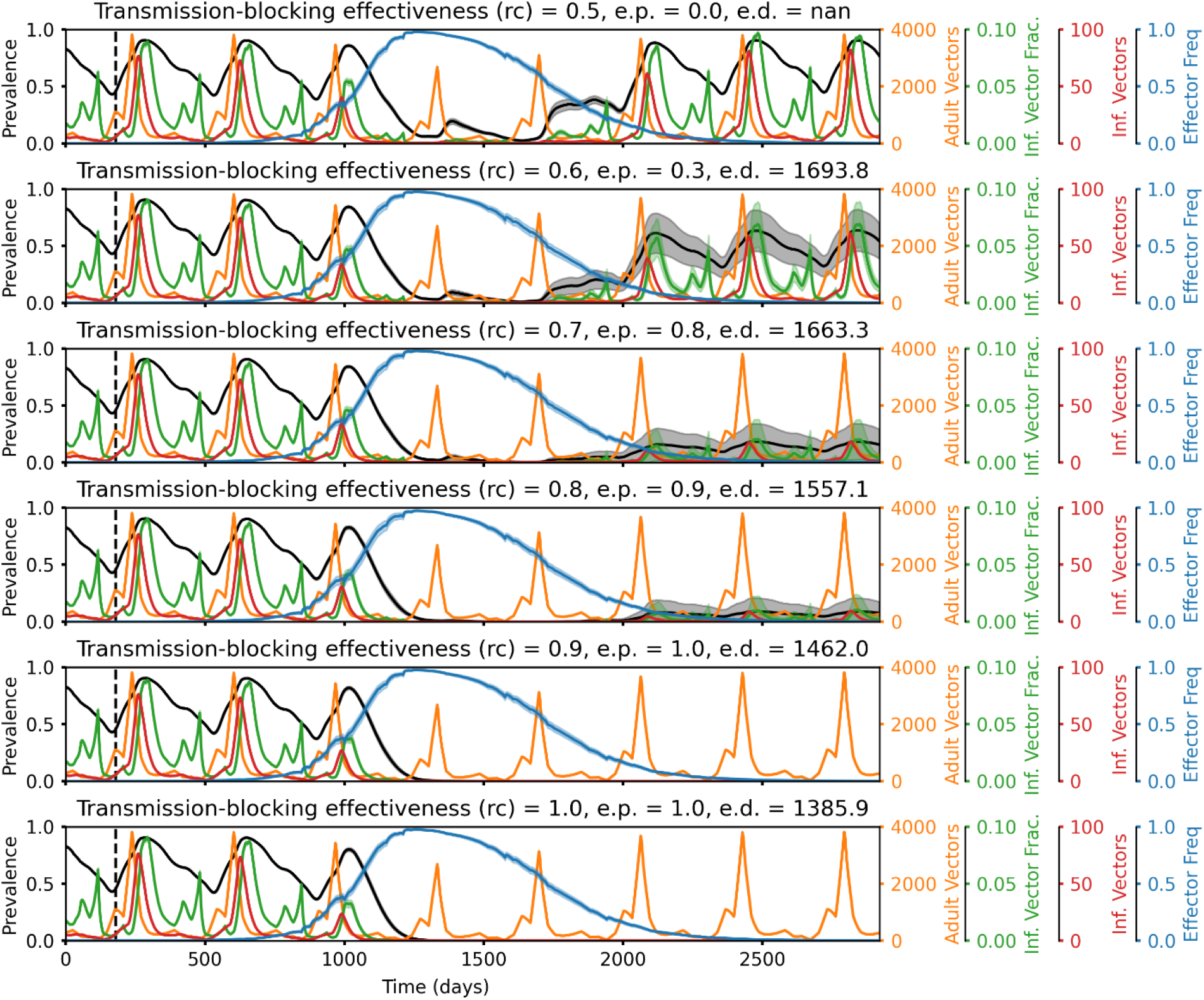
Representative time series illustrating how elimination probabilities increase with increasing transmission-blocking effectiveness. Time series of malaria prevalence, total adult vector population, infectious vector fraction, total infectious adult vector population, and adult vector effector frequency over increasing values of transmission-blocking effectiveness (*rc*). Elimination probabilities (e.p.) and number of days to elimination (e.d.) are denoted in the subplot titles. In the simulations corresponding to these time series, classic gene drive mosquitoes were released in a moderate transmission setting (annual EIR = 30) with non-*rc* parameters set equal to the following values: drive efficiency (*d*) = 1, pre-existing resistance (*rr0*) = 0.01, and fitness cost (*sne*) = 0.4. The higher the transmission-blocking effectiveness, the lower the frequency of vectors that are infectious among the total vector population and the greater the chance of locally eliminating malaria.

**Figure 5.**
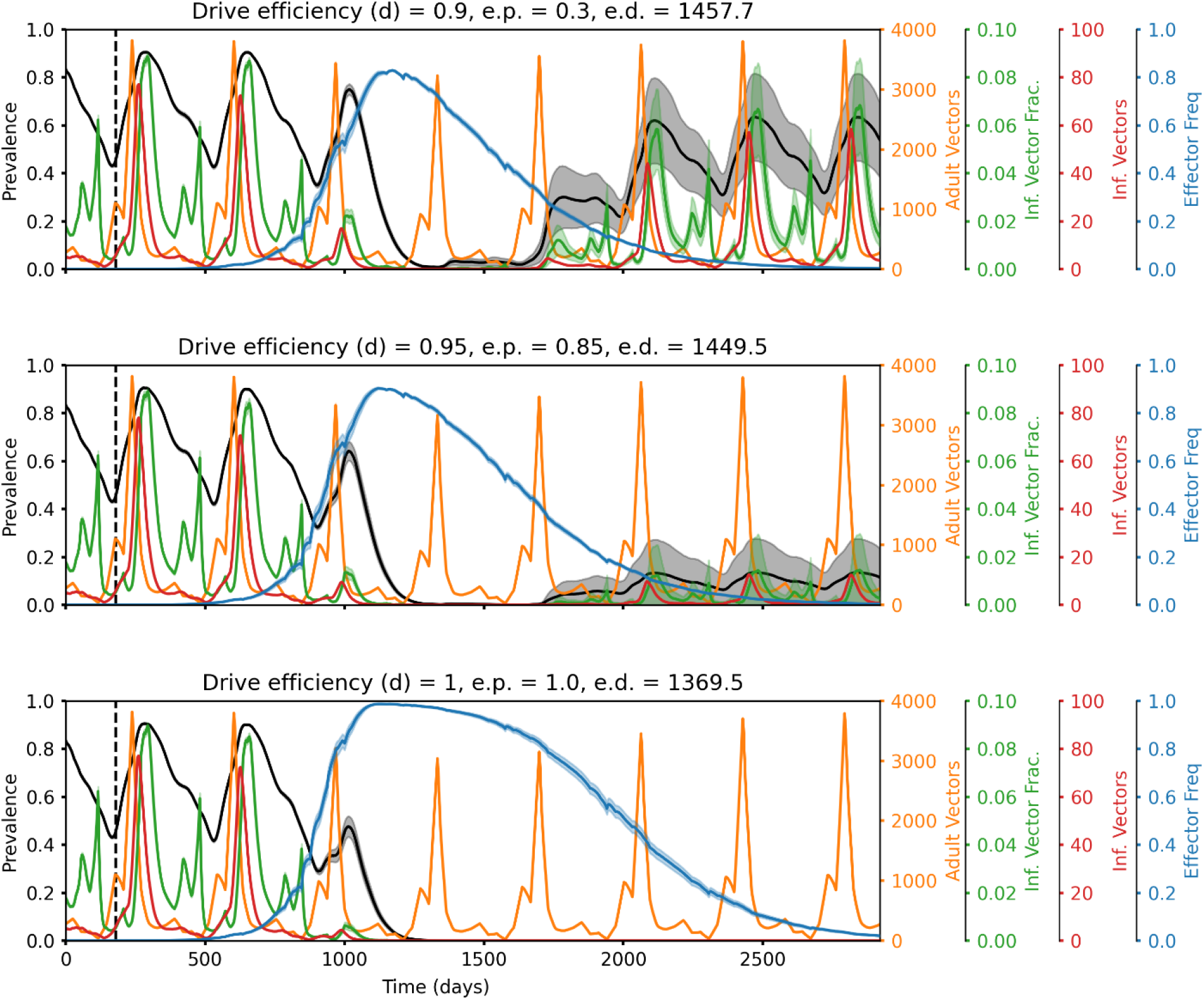
Representative time series illustrating how elimination probabilities increase with increasing drive efficiency. Time series of malaria prevalence, total adult vector population, infectious vector fraction, total infectious adult vector population, and adult vector effector frequency over increasing values of drive efficiency (*d*). Elimination probabilities (e.p.) and number of days to elimination (e.d.) are denoted in the subplot titles. In the simulations corresponding to these time series, classic gene drive mosquitoes were released in a moderate transmission setting (annual EIR = 30) with non-d parameters set equal to the following values: transmission-blocking effectiveness (*rc*) = 0.9, pre-existing resistance (*rr0*) = 0.01, and fitness cost (*sne*) = 0.3. The higher the drive efficiency, the greater the peak effector frequency, the lower the infectious vector fraction, and the greater the chance of locally eliminating malaria.

**Figure 6.**
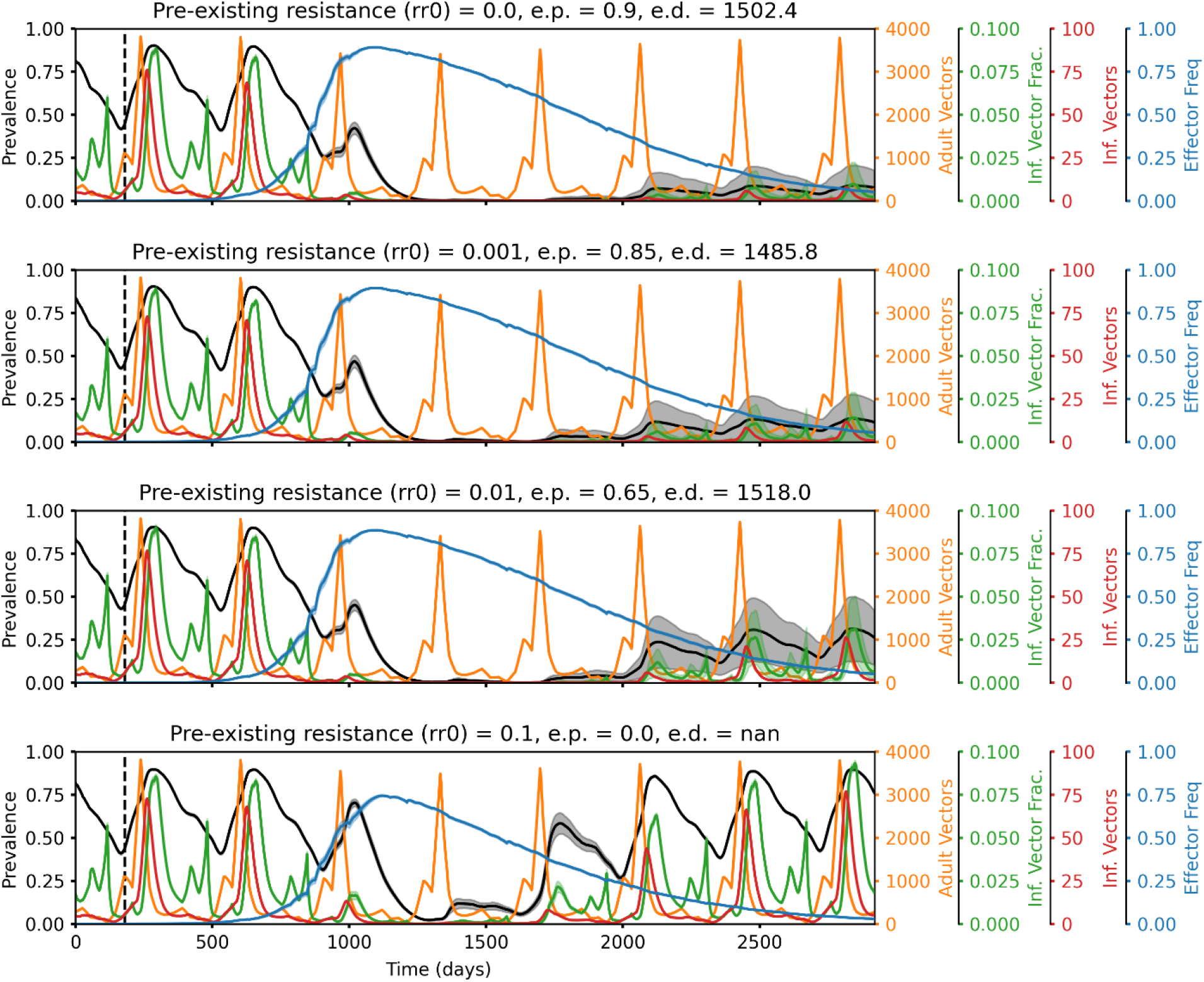
Representative time series illustrating how elimination probabilities decrease with increasing pre-existing population target site resistance. Time series of malaria prevalence, total adult vector population, infectious vector fraction, total infectious adult vector population, and adult vector effector frequency over increasing values of pre-existing population target site resistance frequency (*rr0*). Elimination probabilities (e.p.) and number of days to elimination (e.d.) are denoted in the subplot titles. In the simulations corresponding to these time series, classic gene drive mosquitoes were released in a moderate transmission setting (annual EIR = 30) with non-*rr0* parameters set equal to the following values: drive efficiency (*d*) = 0.9, transmission-blocking effectiveness (*rc*) = 0.9, and fitness cost (*sne*) = 0.2. The higher the pre-existing resistance, the lower the peak effector frequency, the higher the infectious vector fraction, and the lower the chance of locally eliminating malaria.

In comparison to the above three parameters (*rc*, *d*, and *rr0*), the effects of mortality-enhancing fitness costs (*sne*) associated with expression of the introduced gene drive construct have a more complex relationship with the likelihood of elimination. Depending on the context, elimination probabilities can either increase or decrease with increases in fitness cost. In some scenarios, an increase in fitness cost leads to a decrease in elimination probability (Figure 3, columns with upward triangles). Here an increase in fitness cost associated with construct expression both delays and reduces peak effector frequency (Supp. Figure 3A). As fitness costs of expressing the complete construct increase, wild type mosquitoes can more readily outcompete mosquitoes bearing the complete construct, slowing the initial spread of the effector through the population. Resistant mosquitoes can also more readily outcompete mosquitoes with the complete construct at higher fitness costs, such that resistant alleles can increase more rapidly and to a higher frequency in comparison to effector alleles in these situations. These two effects work together to reduce effector frequency at all times in the population and therefore lead to lower elimination probabilities with higher fitness costs. In other scenarios, however, an increase in fitness cost leads to an increase in elimination probability (Figure 3, columns with downward triangles). Here an increase in fitness cost still reduces effector frequency as before; however, a transient reduction in total vector population due to higher mortality rates with higher fitness costs has a larger effect on reducing malaria prevalence than a decrease in effector frequency has on increasing malaria prevalence (Supp. Figure 3B). It is also possible for a combination of the above two situations to occur, such that elimination probabilities can first increase and then decrease with fitness cost (Supp. Figure 3C). In this case, an increase in fitness cost from low initial values substantially reduces the total vector population without significantly affecting effector frequencies; then an increase in fitness cost at moderate to high initial values substantially lowers effector frequencies while only somewhat reducing the total vector population in comparison. Notably, the first scenario (a decrease in elimination probabilities with increasing fitness cost) typically occurs at higher transmission-blocking effectiveness values, while the latter scenario (an increase in elimination probabilities with increasing fitness cost) typically occurs at lower transmission-blocking effectiveness values (Figure 3). This is likely because adult vector numbers matter more and effector frequencies matter less at lower transmission-blocking effectiveness values, since the effector is already relatively pervious. On the other hand, at higher values of transmission-blocking effectiveness, decreases in effector frequency are more detrimental to malaria suppression, as elimination is more dependent on the effector working well. Importantly, all simulations assume that the target species *Anopheles gambiae* is the sole malaria vector and no expansion of other malaria-transmitting species to fill the ecological niche left by a transient decrease in the original number of vectors.

For simulations that result in elimination, trends in elimination timing (defined as the number of simulated years required to reach elimination starting from simulation day 0) follow those of elimination probability (Figure 7). That is, higher elimination probabilities are associated with faster times to elimination and are driven in the same ways by the four tested parameters. Increasing drive efficiency and transmission-blocking effectiveness reduce time to elimination; increasing pre-existing resistance increases time to elimination; and increasing fitness cost can increase or decrease time to elimination depending on the same factors described for elimination probability above. This association between elimination probability and timing occurs because gene drive parameter spaces leading to higher and longer-lasting peak effector frequencies also tend to lead to earlier peak effector frequencies as well.

**Figure 7.**
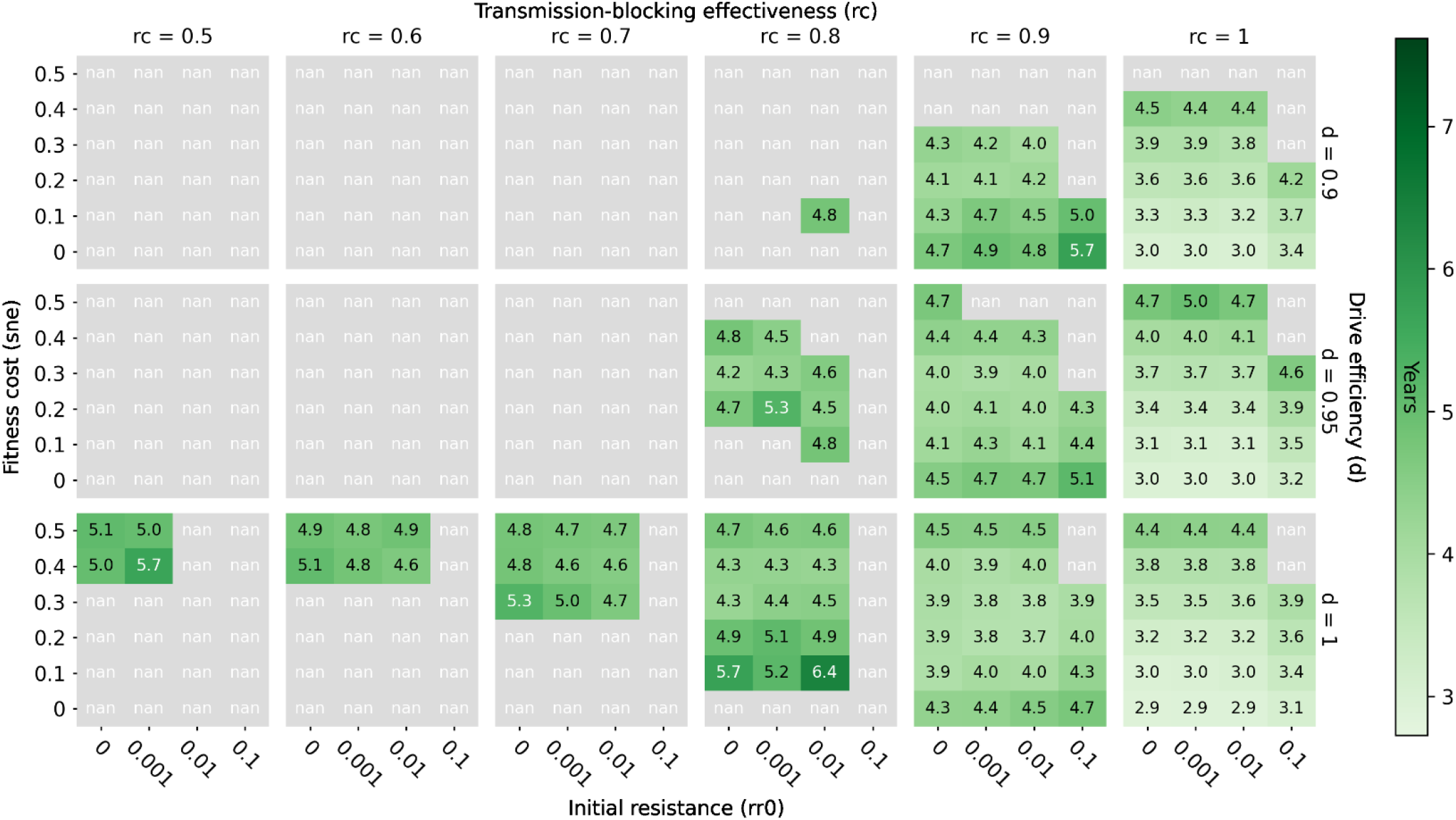
Elimination timing after a single release of classic gene drive mosquitoes only in a moderate transmission (annual EIR = 30) regime. Elimination timing (computed as the number of years taken to reach elimination starting from simulation day 0, averaged over all realizations that eliminate) over a range of transmission-blocking effectiveness (*rc*), drive efficiency (*d*), pre-existing population target site resistance frequency (*rr0*), and mortality-enhancing effector expression fitness cost (*sne*) values.

The above-described patterns of elimination probability driven by transmission-blocking effectiveness, drive efficiency, pre-existing target site resistance, and fitness cost associated with a single classic gene drive mosquito release also hold when integral rather than classic gene drive mosquitoes are released, as well as when ITNs are deployed in addition to a single release of either classic or integral gene drive mosquitoes (Figures 8–9). In the case of additional ITN deployment, however, increases in drive efficiency do not always lead to increases in elimination probability because of mismatches in seasonality and timing of maximum net and gene drive efficacy within the setups simulated here (gene drive mosquitoes released on July 1 of year 1 and ITNs deployed by July 1 of year 1, 4 and 7). As was the case for gene drive only scenarios, when drive efficiency increases, the peak in effector frequency shifts earlier. In some situations with additional ITN deployment, however, this earlier peak in effector frequency then subsides by the time ITNs are re-deployed a second time (on July 1 of year 4 - after waning efficacy of ITNs from the initial deployment), such that the overlapping maximum effects of ITNs and gene drive mosquitoes during low season (when chances of eliminating are highest) are actually smaller than if drive efficiency were lower and peak effector frequency were more delayed (Supp. Figure 4). Timing the release of gene drive mosquitoes such that peak effector frequency coincides with maximum ITN efficacy and the smallest vector population size may therefore be key to eliminating malaria in certain situations.

**Figure 8.**
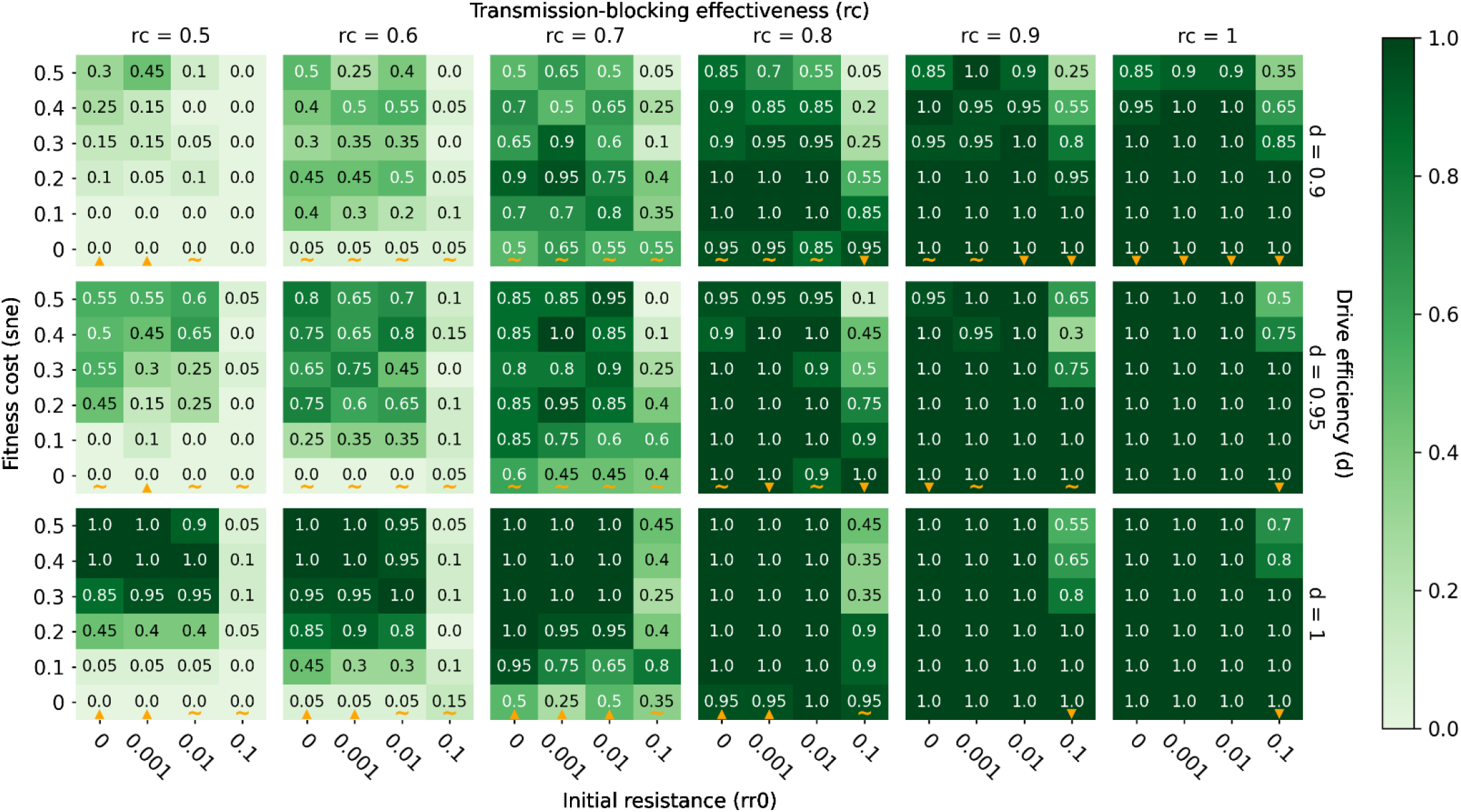
Elimination probabilities after a single release of classic gene drive mosquitoes and ITN deployment in a moderate transmission (annual EIR = 30) regime. Same as Figure 3.

**Figure 9.**
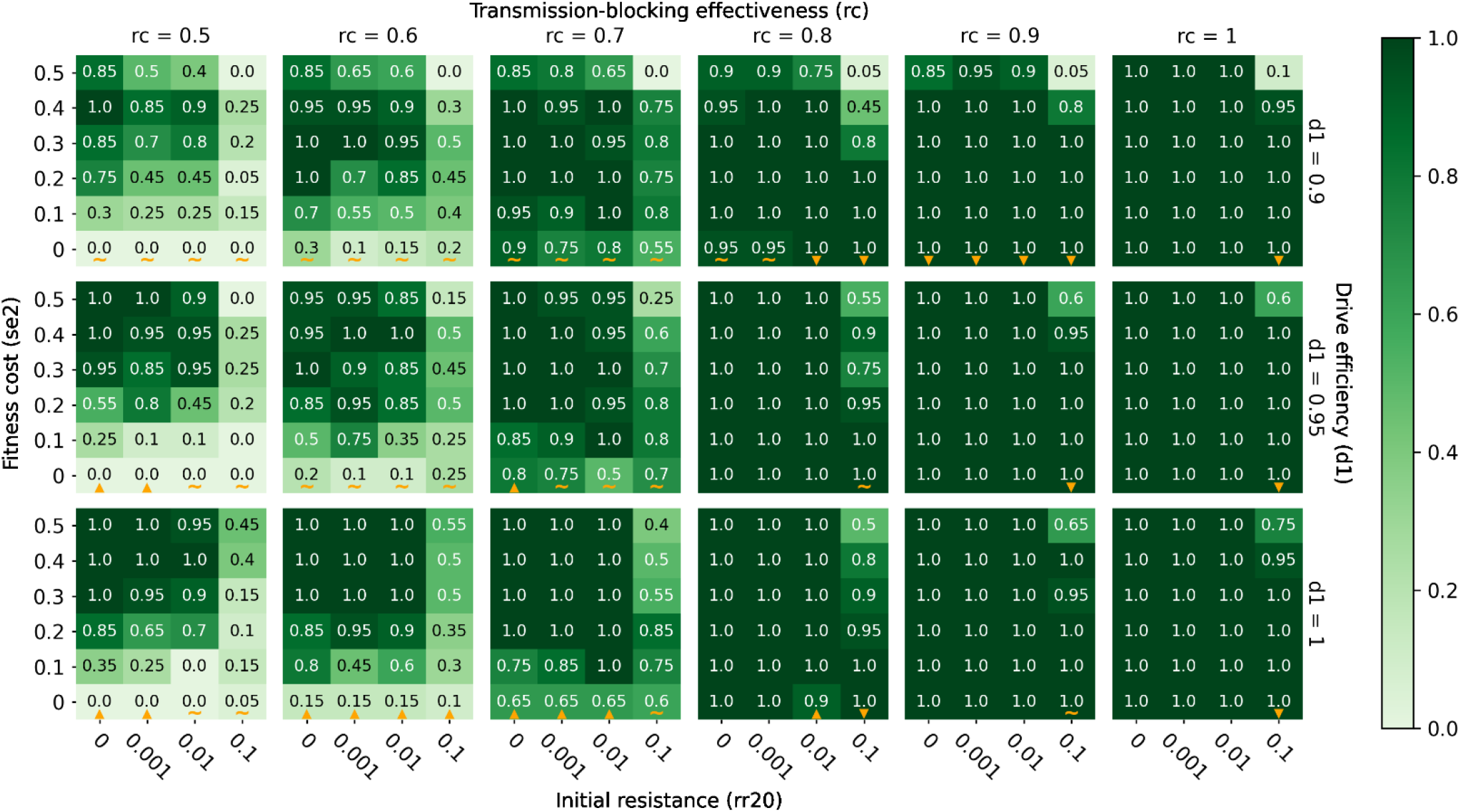
Elimination probabilities after a single release of integral gene drive mosquitoes and ITN deployment in a moderate transmission (annual EIR = 30) regime. Same as Figure 3.

### A single release of classic gene drive mosquitoes with high transmission-blocking effectiveness is likely to locally eliminate malaria in low to moderate transmission settings

A single release of classic gene drive mosquitoes can virtually guarantee elimination in a moderate transmission regime (annual EIR = 30) if the transmission-blocking effectiveness is greater than or equal to 90%, pre-existing target site resistance is less than or equal to 1%, drive efficiency is greater than or equal to 95%, and the fitness cost to mortality of expressing the effector is less than or equal to 40% (Figure 3). If transmission-blocking effectiveness is 100% and drive efficiency is greater than or equal to 95%, pre-existing target site resistance can be as high as 10% if fitness costs are below ^~^20% (Figure 3). When drive efficiency is less than or equal to 95%, a transmission-blocking effectiveness of ^~^80% or less makes elimination highly unlikely or virtually impossible (Figure 3). In a low transmission regime (annual EIR = 10), elimination probabilities are appreciably higher across all parameter values, such that a transmission-blocking effectiveness of 80% (rather than 90%) still leads to high probabilities of elimination even when drive efficiency is less than or equal to 95% (Supp. Figure 5).

### Deployment of ITNs in conjunction with a single classic gene drive mosquito release boosts elimination probability

Although a single release of classic gene drive mosquitoes with high transmission-blocking effectiveness is likely to locally eliminate malaria by itself in low to moderate transmission regimes, additional deployment of ITNs greatly enhances elimination probabilities at lower values of transmission-blocking effectiveness (Figure 8). By deploying ITNs in a moderate transmission regime (annual EIR = 30), elimination goes from virtually impossible to highly likely when transmission-blocking effectiveness drops down to ^~^70% and drive efficiency is less than or equal to 95% (Figure 8). In the absence of ITNs, elimination probabilities are negligible when transmission-blocking effectiveness is less than or equal to 80% and drive efficiency is less than or equal to 95% (Figure 3). In a low transmission regime (annual EIR = 10), additional ITN deployment leads to high probabilities of elimination at values of transmission-blocking effectiveness as low as 50% (Supp. Figure 6). In a high transmission regime (annual EIR = 80), ITN deployment and classic gene drive mosquito release in combination lead to high probabilities of elimination when transmission-blocking effectiveness is ^~^80% or higher (Supp. Figure 7).

### Release of integral, rather than classic, gene drive mosquitoes further boosts elimination probabilities

In comparison to a release of classic gene drive mosquitoes, a release of integral gene drive mosquitoes expands the parameter space over which elimination is highly likely (Figure 9). That is, an integral gene drive mosquito release can achieve the same or better elimination outcomes as a classic gene drive mosquito release even with a less potent effector, a lower drive efficiency, and in the presence of a higher target site resistance frequency (here at the effector site) within the population. In a moderate transmission regime (annual EIR = 30), releasing integral gene drive mosquitoes in conjunction with ITN deployment leads to high and near certain elimination probabilities at transmission-blocking effectiveness values as low as 50% and drive efficiencies as low as 90% (Figure 9), compared to transmission-blocking effectiveness values around 70% and similar drive efficiencies when releasing classic gene drive mosquitoes (Figure 8). An integral gene drive mosquito release in low and high transmission regimes yields similar increases in elimination probabilities compared to a classic gene drive mosquito release (Supp. Figures 8–9). To visualize elimination probabilities (along with elimination timing, prevalence, vector population, and allele frequencies) for all simulated combinations of gene drive release types (classic and integral), ITN deployments (with and without), and transmission regimes (annual EIR = 10, 30, and 80), see https://gene-drive.bmgf.io.

## Discussion

### Transmission-blocking effectiveness and fitness cost may be the most important parameters for future improvements and characterizations

Transmission-blocking effectiveness, drive efficiency, pre-existing target site resistance, and fitness cost of expressing an introduced anti-pathogenic effector were all important parameters affecting local malaria elimination probabilities across all simulated transmission intensities and scenarios. In general, elimination probabilities were highest when transmission-blocking effectiveness was highest, drive efficiency was highest, and pre-existing target site resistance was lowest (Figures 3, 8–9; Supp. Figures 5–9). When deploying ITNs together with a gene drive release, however, increased drive efficiencies did not always increase elimination probabilities due to mismatches in timing between maximum ITN efficacy and peak effector frequency. To increase the chances of elimination, it is therefore important to accurately quantify the genetic parameters associated with a given gene drive mosquito strain of interest and time its release such that peak effector frequency coincides with both low mosquito season and maximum efficacy of other forms of traditional vector control. Extensive vector surveillance before gene drive mosquito release would also be needed to accurately quantify vector population seasonality, along with pre-existing target site resistance.

Fitness cost affected elimination probability differently depending on transmission-blocking effectiveness. At very high values of transmission-blocking effectiveness, higher fitness costs reduced elimination probabilities, while at lower values of transmission-blocking effectiveness, this effect was reversed (Figures 3, 8–9; Supp. Figures 5–9). Because fitness cost effects on elimination probabilities are not uniform or easily predicted, researchers and public health workers should develop a good understanding of both transmission-blocking effectiveness and fitness costs associated with their gene drive mosquito strains of interest before release. Semi-field experiments and non-driving effector releases could be instrumental in achieving this goal, as the translation of experimentally established fitness parameters into actual fitness burden incurred by transgenic mosquitoes in the environment is notoriously difficult. Extensive vector surveillance should also be conducted after all gene drive mosquito releases, but especially for pilot releases, to validate models and better understand complicated effects of gene drive system parameters such as fitness cost. Sufficient surveillance after release can also be used to track failure rates and inform necessary adjustments to future gene drive strains or release logistics.

Though all four parameters tested here had some measurable effect on elimination probabilities, transmission-blocking effectiveness and fitness cost may be most important to focus on for future improvements to new strains of gene drive mosquitoes, due both to their outsized influence on elimination probability as well as their potentially limiting existing values.

#### Pre-existing target site resistance

Existing pre-existing target site resistances in wild *An. gambiae* populations will likely not adversely affect the ability of population-replacing classic or integral gene drive mosquito releases to eliminate malaria. Using a sample of ^~^1,000 wild-caught *An. gambiae s.l*. mosquitoes from natural populations throughout Africa, Schmidt et al. (2020) [50] found that the vast majority (^~^90%) of all protein-coding genes in *An. gambiae s.l*. contain at least one Cas9 target sequence with genetic variability less than or equal to 1%. This sample of ^~^1,000 mosquitoes included both *An. coluzzii* and *An. gambiae s.s*. specimens from the UC Davis Vector Genetics Laboratory and The *Anopheles gambiae* 1000 Genomes Consortium. Furthermore, though genetic variability may be present, even when target sequences differ by multiple nucleotides, efficient cleavage by a Cas9 driver enzyme may remain largely unimpaired [51,52]. Existing population target site resistances are therefore likely to be low (less than or equal to 1%), given the ability of researchers to choose a favorable site with little Cas9-impairing genetic variability.

#### Drive efficiency

Given realistic values of ^~^90-100% for drive efficiencies in *Anopheles* mosquitoes [51–55], elimination probabilities are not substantially reduced when drive efficiency decreases within this range and transmission-blocking effectiveness is sufficiently high (e.g., greater than or equal to ^~^90% when releasing classic gene drive mosquitoes without ITNs in a transmission regime where annual EIR = 30, and above ^~^60-70% in the same situation with ITNs). This is true for releases of classic or integral gene drive mosquitoes, with or without vector control, though the exact values of transmission-blocking effectiveness required differ depending on the gene drive system, transmission regime, and absence or presence of other forms of vector control. Thus, even at the lower end of realistic *Anopheles* drive efficiency values (^~^90%), elimination probabilities are generally not limited by drive transmission rates. Though increasing drive efficiencies from 95% to 100% can boost elimination probabilities at lower transmission-blocking effectiveness values (along with high fitness costs and low but realistic pre-existing target site resistances), drive efficiencies of ^~^95% may be extremely difficult to improve upon with conventional mosquito engineering efforts. Thus, drive transmission rate is not a high priority for further improvements due to its already high efficiency and promising ability to enable elimination, along with the likely difficulty associated with bringing efficiencies even higher.

#### Fitness cost

Fitness costs associated with expressing an anti-parasite effector have been reported to vary widely among engineered strains of *An. gambiae* under laboratory conditions. Some anti-parasite effector expressing strains have been created with negligible associated fitness costs [56,57], while others exhibit measurable potential decreases in fecundity or lifespan [57,58]. It would be theoretically favorable to create strains with effector expression fitness costs as low as possible, since lower fitness costs allow introduced anti-pathogenic GM strains to more readily spread and compete against wild type mosquitoes. However, fitness cost ranges required to achieve elimination are highly dependent on other parameters. Assuming the presence of one primary malaria vector species and limited niche expansion by another, a single release of gene drive mosquitoes with lower transmission-blocking effectiveness is more likely to eliminate malaria when associated fitness costs of effector expression are higher (up to a certain point). On the other hand, a release of more effective transmission-blocking gene drive mosquitoes may be increasingly likely to eliminate malaria at lower fitness costs. Thus, rather than universally seeking to generate strains with reduced fitness costs, researchers may opt to generate strains with optimal combinations of drive efficiency, fitness cost, and transmission-blocking effectiveness to increase the chances of elimination in their particular setting of interest. Though transmission-blocking effectiveness must always be above some minimum threshold for any population replacement gene drive release to achieve elimination, there is no equivalent maximum threshold that fitness cost must be below. Here we simulated fitness cost as a uniform increase in vector mortality across all ages and sexes, but future work could examine the outcomes of age or sex-specific fitness effects, such as a reduction in the lifespan of females only.

#### Transmission-blocking effectiveness

Transgenic strains of *An. stephensi* with *anti-falciparum* transmission-blocking effectivenesses of 100% or nearly 100% have been created [20,22,59,60]. However, there does not yet exist a transgenic strain of *An. gambiae* that is able to inhibit *P. falciparum* parasite transmission as completely. Indeed, most transgenic *An. gambiae* effectors show modest reductions in parasite transmission ability [57,58,61,62]. The most effective transgenic *An. gambiae* strain we were able to find in the published literature was able to reduce the number of sporozoites per salivary gland by ^~^50% [57]. In the case of a single integral gene drive mosquito release with ITNs in a high transmission setting (annual EIR = 80), elimination probabilities drop precipitously below transmission-blocking effectiveness values of 60-70%, assuming realistic ranges of other parameters. Thus, existing transmission-blocking effectivenesses in *An. gambiae* are generally unlikely to be high enough to achieve elimination in high transmission settings even when gene drive mosquito release is deployed in combination with ITNs. Transmission-blocking effectiveness should therefore be the most important primary focus for future improvements in new strains of gene drive mosquitoes.

### Existing population replacement gene drive mosquitoes, in combination with traditional forms of vector control, can likely eliminate malaria in low to moderate transmission settings

A single release of a few hundred highly effective anti-pathogen, population-replacing classic gene drive mosquitoes can locally eliminate malaria in a highly seasonal Sahelian setting with moderate transmission rates (annual EIR of 30) when transmission-blocking effectiveness is very high (^~^90% or higher) and other parameters (drive efficiency, pre-existing target site resistance, and fitness cost of effector expression) are within realistic ranges (Figure 3). When paired with ITN deployment, a single release of a few hundred classic gene drive mosquitoes increases the probability of elimination at all values of drive efficiency, pre-existing target site resistance, transmission-blocking effectiveness, and fitness cost (Figure 9). With ITNs, elimination probabilities are substantial even at transmission-blocking effectiveness values down to ^~^50%. Thus, pairing a population replacement gene drive release with provision of ITNs enhances rather than reduces the effect of the gene drive release, as was also shown by Selvaraj et al. (2020) [32]. Utilizing integral gene drive mosquitoes with separate driver and effector genes inserted at different loci also increases elimination probabilities across the board (Figure 9). In a low transmission regime (annual EIR of 10), a single release of a few hundred classic gene drive mosquitoes can eliminate malaria when transmission-blocking effectiveness is again very high, although less so (^~^80% or higher) (Supp. Figure 5). To ensure high probabilities of elimination at lower transmission-blocking effectiveness values (down to ^~^50%), integral gene drive mosquitoes, along with ITNs and/or other forms of vector control, should again be utilized (Supp. Figure 6, 8). Thus, ITNs and other traditional, non-gene drive vector control strategies are essential tools in the path towards elimination because a single release of mosquitoes with currently achievable gene drive characteristics is not likely to achieve elimination on its own, even in a low transmission regime. In a high transmission setting (annual EIR = 80), additional non-gene drive interventions become even more important. In this regime, transmission-blocking effectiveness values of ^~^50% lead to high probabilities of elimination only when fitness costs are within a narrow range. Therefore, elimination in a high transmission Sahelian setting will likely require a vast improvement to transmission-blocking effectiveness in future integral gene drive mosquito strains and/or additional layering of non-gene drive interventions beyond ITNs. These other interventions could include short term strategies such as indoor residual spraying (IRS), attractive targeted sugar baits (ATSBs), long-acting injectable anti-malarials, and larviciding, as well as longer term approaches including housing improvement, environmental management, and health systems strengthening. Regardless of which other interventions are utilized, releasing population-replacing gene drive mosquitoes helps create a window of opportunity during which prevalence may be greatly suppressed and other tools can be ramped up to achieve elimination, even when a single gene drive mosquito release by itself cannot.

### Model limitations and future work

We sought to present as comprehensive and accurate an overview as possible of the effects of a single gene drive mosquito release on malaria elimination within a spatially resolved and realistically seasonal Sahelian setting. However, we made many necessary simplifying assumptions and were not able to address all possible potentially relevant factors in this initial study. First, though releasing a larger number of mosquitoes within the same 6 most populous nodes did not significantly affect our results, we did not test whether altering the spatial pattern of mosquito release would measurably alter elimination probabilities. Future work can optimize gene drive mosquito release locations and examine factors contributing to optimal spatial planning for releases. Second, evolution and development of parasite resistance to the anti-parasite effector molecules within the mosquitoes is not captured in our model [63]. Future work can seek to better understand how this type of evolution could affect both timing and probability of elimination probabilities, though parasite resistance can also be mitigated by releasing a second set of mosquitoes with a different type of effector that would be new to the parasite. Third, we only accounted for one species and pool of mosquitoes (*Anopheles gambiae*) in our simulations and assumed that other species were either not present or did not play an appreciable role in malaria transmission. If another malaria-transmitting *Anopheles* species were present to fill the ecological niche of the single simulated species, elimination probabilities likely would not increase as substantially with higher fitness costs and reduced vector populations at low values of transmission-blocking effectiveness. Future work can examine elimination probabilities in the presence of multiple malaria-transmitting *Anopheles* species with releases of gene drive mosquitoes corresponding to each different species. Future work can also explore the effects of multiple gene drive mosquito releases over several years compared to a one-time release. Lastly and perhaps most importantly, though we simulated human and vector migration between 1 km-by-1 km nodes within the region, we did not include migration of humans or vectors into or out of the simulated region. While the inner nodes from the simulations serve as a proxy to study the effects of migration from outside regions to the simulated area, we realize that continued importation of malaria via humans or vectors from outside of the simulated region could have made elimination more difficult to achieve within the region across all scenarios. However, because the gene drive systems simulated here are self-propagating, genes introduced via these systems would gradually become established in surrounding regions as well, spreading into all vector populations of the same species until a barrier to vector migration and therefore gene flow is reached. Most or all vector populations migrating back into the simulated region would therefore eventually have experienced their own introduction of gene drive mosquitoes as well. Thus, it is not inconceivable that importation of malaria via migrating vectors and/or human travelers into this relatively small region would gradually decrease and potentially become negligible over time. In addition to the greatly reduced importation of malaria via gene drive mosquitoes from outside of the simulated region, human importation of malaria into the simulated region could be greatly reduced if, for example, travelers are required to be tested before entering or returning home. Future larger spatial scale (and therefore lower resolution) simulations, along with incorporation of as yet unavailable additional data on both human and vector migration distances, timings, and frequencies would allow us to better resolve these dynamics. This additional data on vector movements would be invaluable for better understanding the potential spatiotemporal evolution of gene drive mosquito frequencies and the resultant effects on malaria transmission in SSA. Immediate future research should therefore prioritize entomological surveillance efforts.

---

**Supp. Figure 1.**
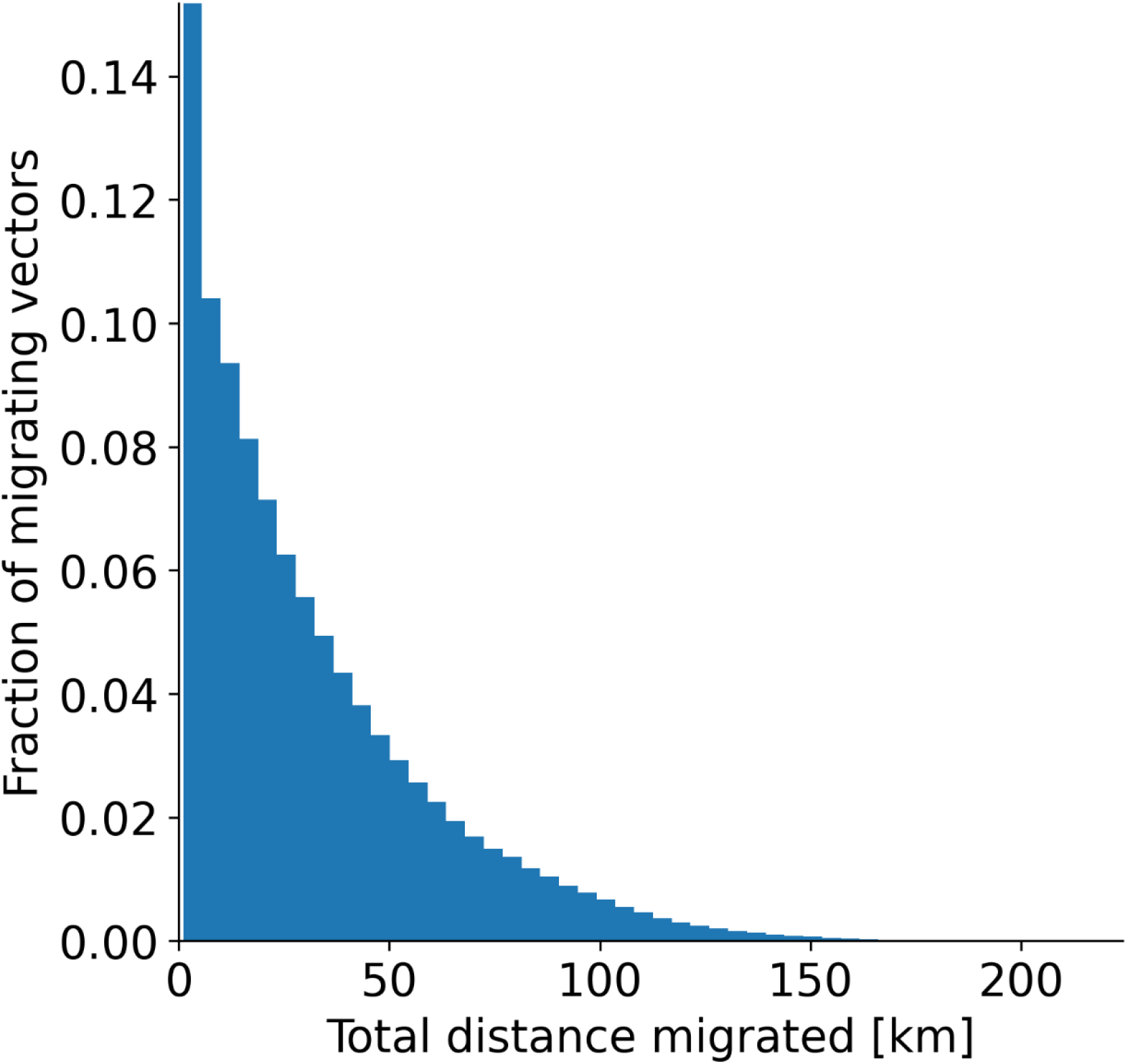
Distribution of vector migration distances within the simulated area. Fraction of vector migrations versus distance, computed by summing total migration distance over each migrating vector’s existence within a 2-month period (August 1 to October 1 in the first simulation year with annual EIR = 30 and no ITNs or gene drive release), counting the number of total migration distances within each histogram distance bin, and then dividing by the total number of migrating vectors in the 2-month period. Total migration distance as plotted here does not necessarily represent the distance between a vector’s starting and ending point (i.e, its displacement), but instead represents the total distance traveled. Migration probabilities are governed by an empirical negative exponential distance decay function [48].

**Supp. Figure 2.**
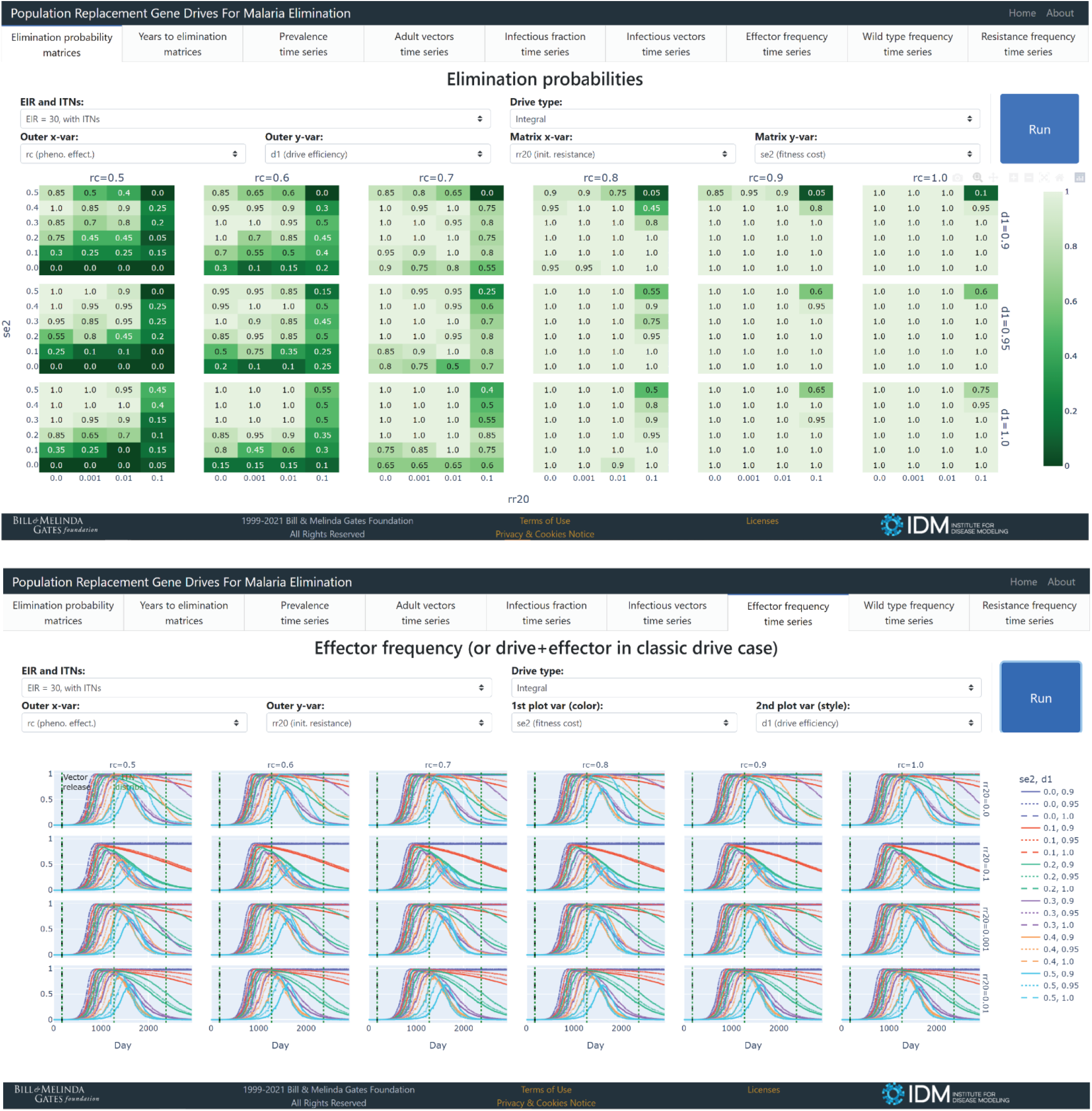
Screenshots of accompanying website for interactive visualization of simulation output. Screenshots of two different tabs on the website located here: https://gene-drive.bmgf.io. Website users can interactively visualize the effects of tested gene drive parameters on elimination probabilities, elimination timing, prevalence, vector populations, and allele frequencies over all simulated combinations of gene drive release types, ITN deployments, and transmission regimes.

**Supp. Figure 3.**
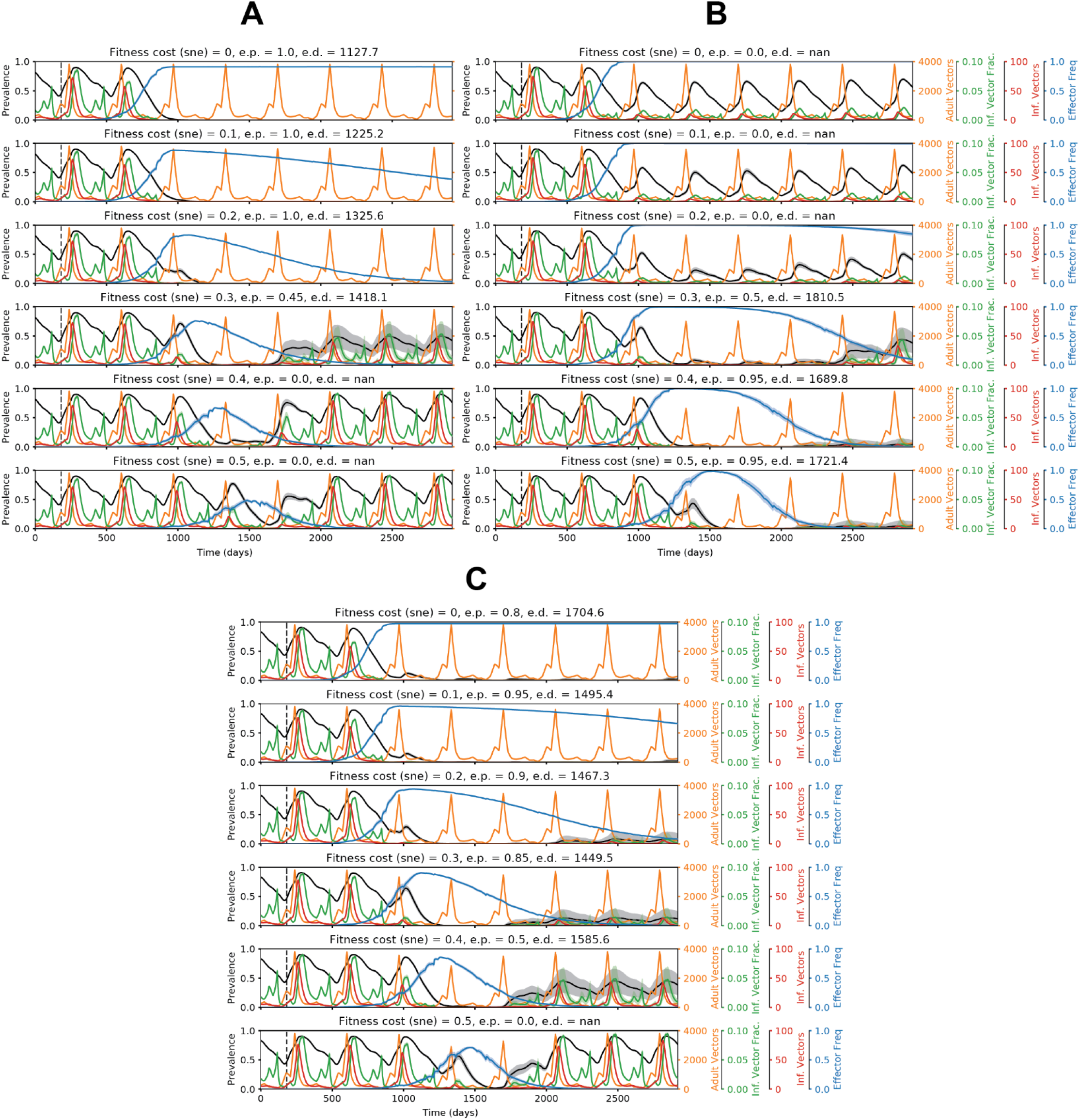
Representative time series illustrating how elimination probabilities can either increase or decrease with increasing fitness costs of complete construct expression. Time series of malaria prevalence, total adult vector population, infectious vector fraction, total infectious adult vector population, and adult vector effector frequency over increasing values of fitness costs associated with complete construct expression (*sne*). Elimination probabilities (e.p.) and number of days to elimination (e.d.) are denoted in the subplot titles. In the simulations corresponding to these time series, classic gene drive mosquitoes were released in a moderate transmission setting (annual EIR = 30). In column A, representing the case in which increasing fitness costs increase elimination probabilities, non-*sne* parameters were set equal to the following values: drive efficiency (*d*) = 1, pre-existing resistance (*rr0*) = 0.001, and transmission-blocking effectiveness (*rc*) = 0.7. In column B, representing the case in which increasing fitness costs decrease elimination probabilities, non-*sne* parameters were set equal to the following values: *d* = 1, *rr0* = 0.1, and *rc* = 1. In column C, representing the case in which increasing fitness costs increase and then decrease elimination probabilities, non-*sne* parameters were set equal to the following values: *d* = 0.95, *rr0* = 0.01, and *rc* = 0.9. In column A, the higher the fitness costs, the lower the total vector population, and the greater the chance of locally eliminating malaria. In column B, the higher the fitness costs, the lower the peak effector frequency, and the lower the chance of elimination. In column C, the effects described for columns A and B are both at play.

**Supp. Figure 4.**
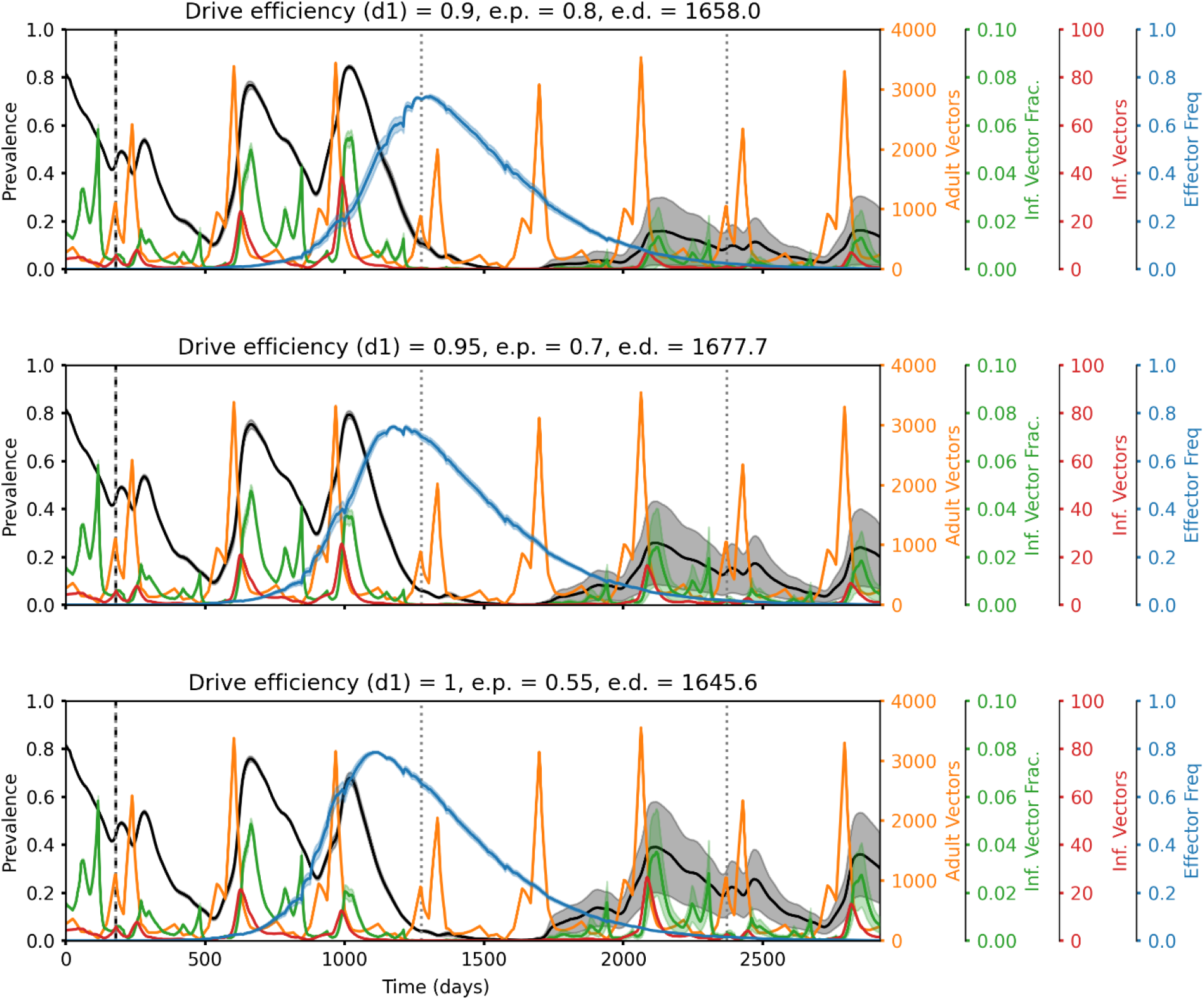
Representative time series illustrating how elimination probabilities can sometimes decrease with increasing drive efficiency. Time series of malaria prevalence, total adult vector population, infectious vector fraction, total infectious adult vector population, and adult vector effector frequency over increasing values of drive efficiency (*d1*). Elimination probabilities (e.p.) and number of days to elimination (e.d.) are denoted in the subplot titles. In the simulations corresponding to these time series, integral gene drive mosquitoes were released and ITNs were deployed in a moderate transmission setting (annual EIR = 30) with non-*d1* parameters set equal to the following values: transmission-blocking effectiveness (*rc*) = 0.7, pre-existing resistance at the effector target site (*rr20*) = 0.1, and fitness cost of expressing the effector (*se2*) = 0.3. The higher the drive efficiency, the earlier the peak in effector frequency, and the lower the chance of locally eliminating malaria when this earlier peak does not match up with maximum ITN efficacy.

**Supp. Figure 5.**
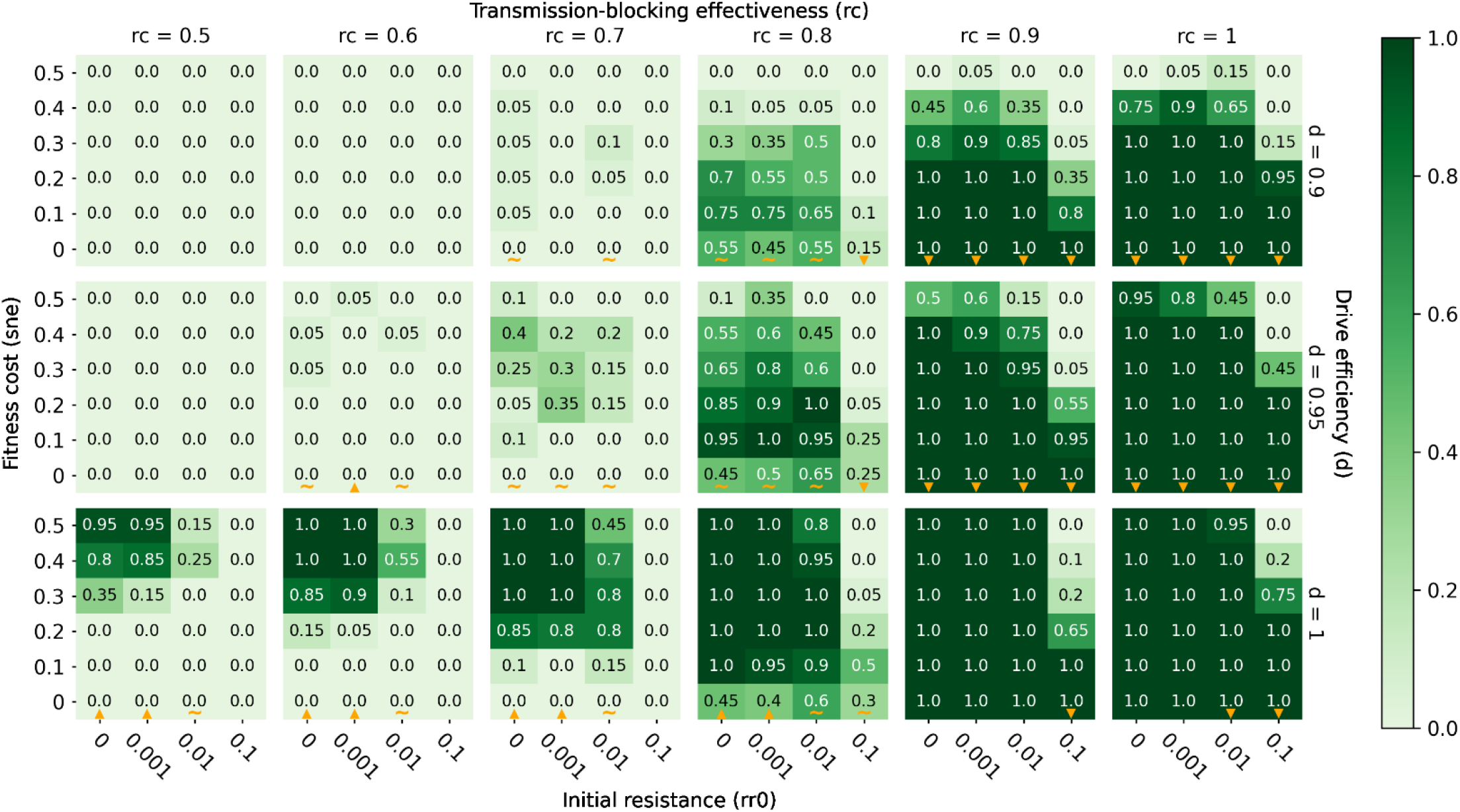
Elimination probabilities after a single release of classic gene drive mosquitoes only in a low transmission (annual EIR = 10) regime. Same as Figure 3.

**Supp. Figure 6.**
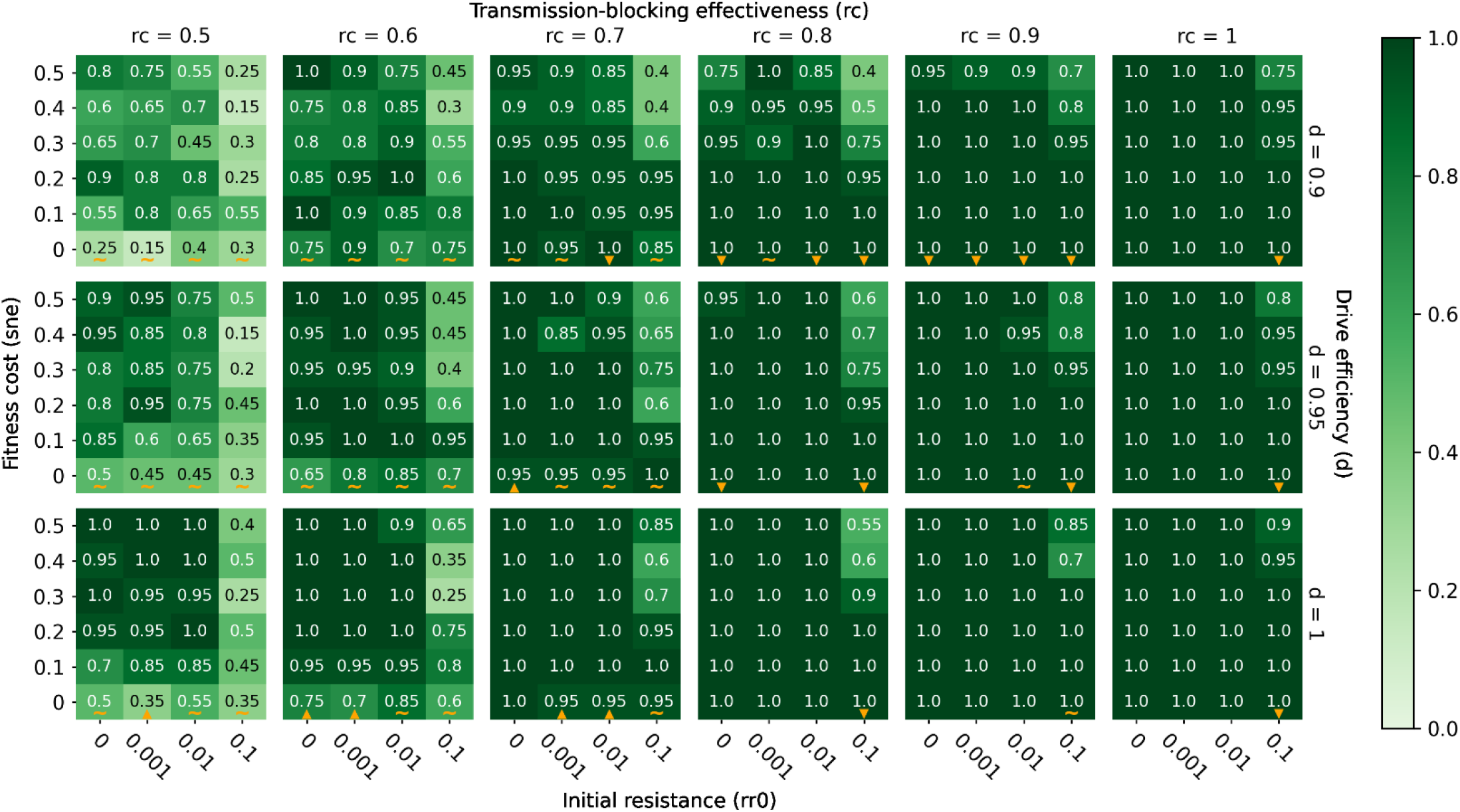
Elimination probabilities after a single release of classic gene drive mosquitoes and ITN deployment in a low transmission (annual EIR = 10) regime. Same as Figure 3.

**Supp. Figure 7.**
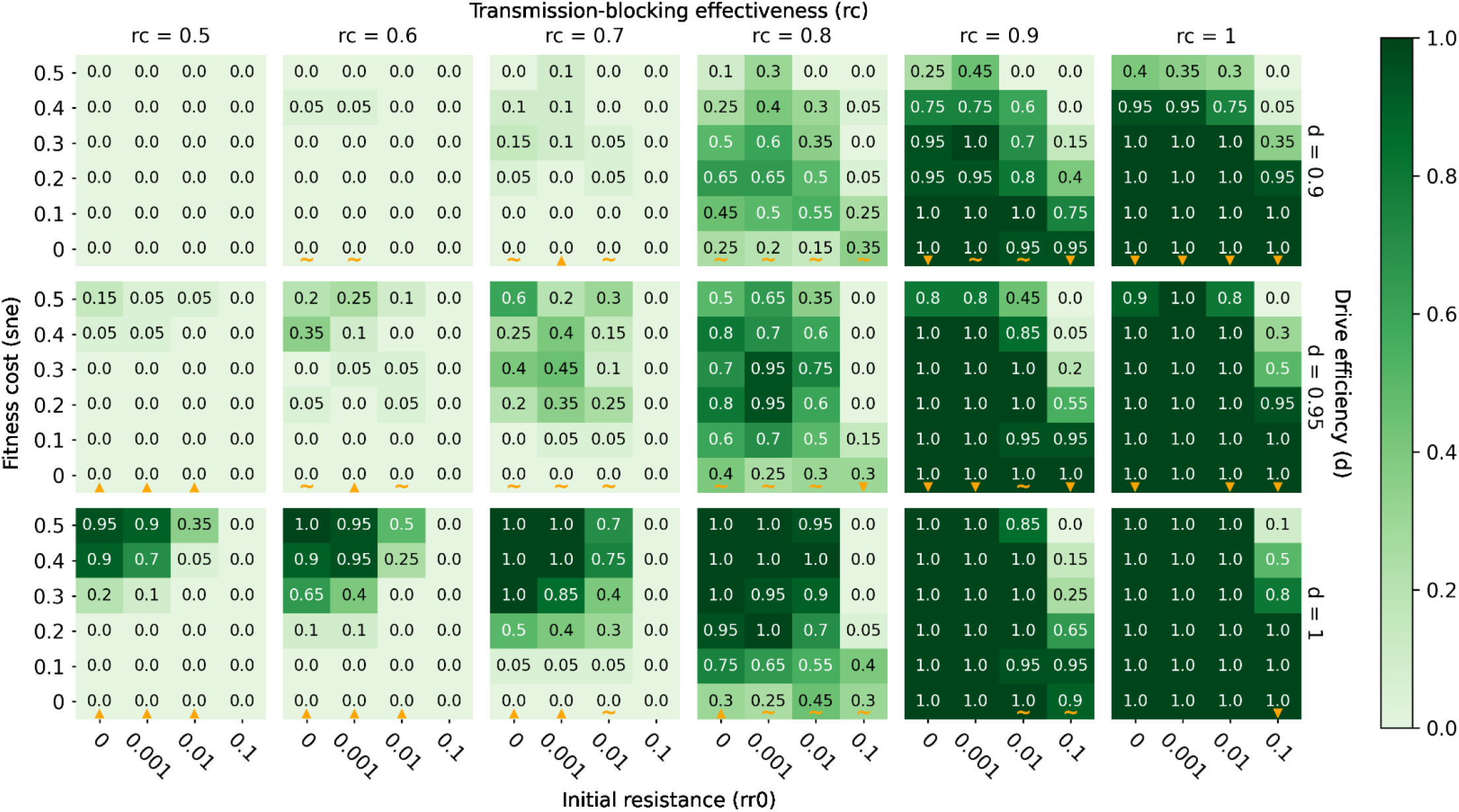
Elimination probabilities after a single release of classic gene drive mosquitoes and ITN deployment in a high transmission (annual EIR = 80) regime. Same as Figure 3.

**Supp. Figure 8.**
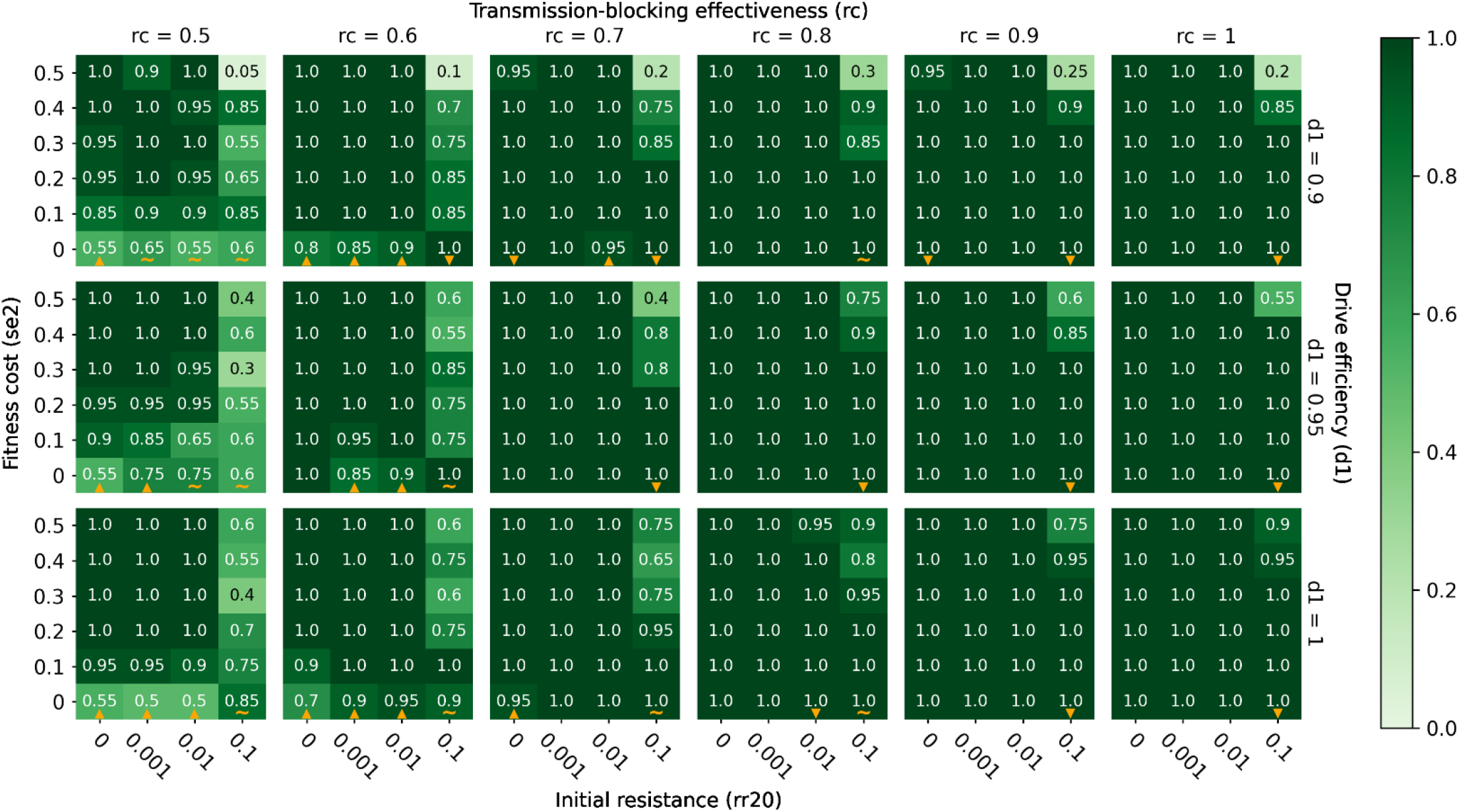
Elimination probabilities after a single release of integral gene drive mosquitoes and ITN deployment in a low transmission (annual EIR = 10) regime. Same as Figure 3.

**Supp. Figure 9.**
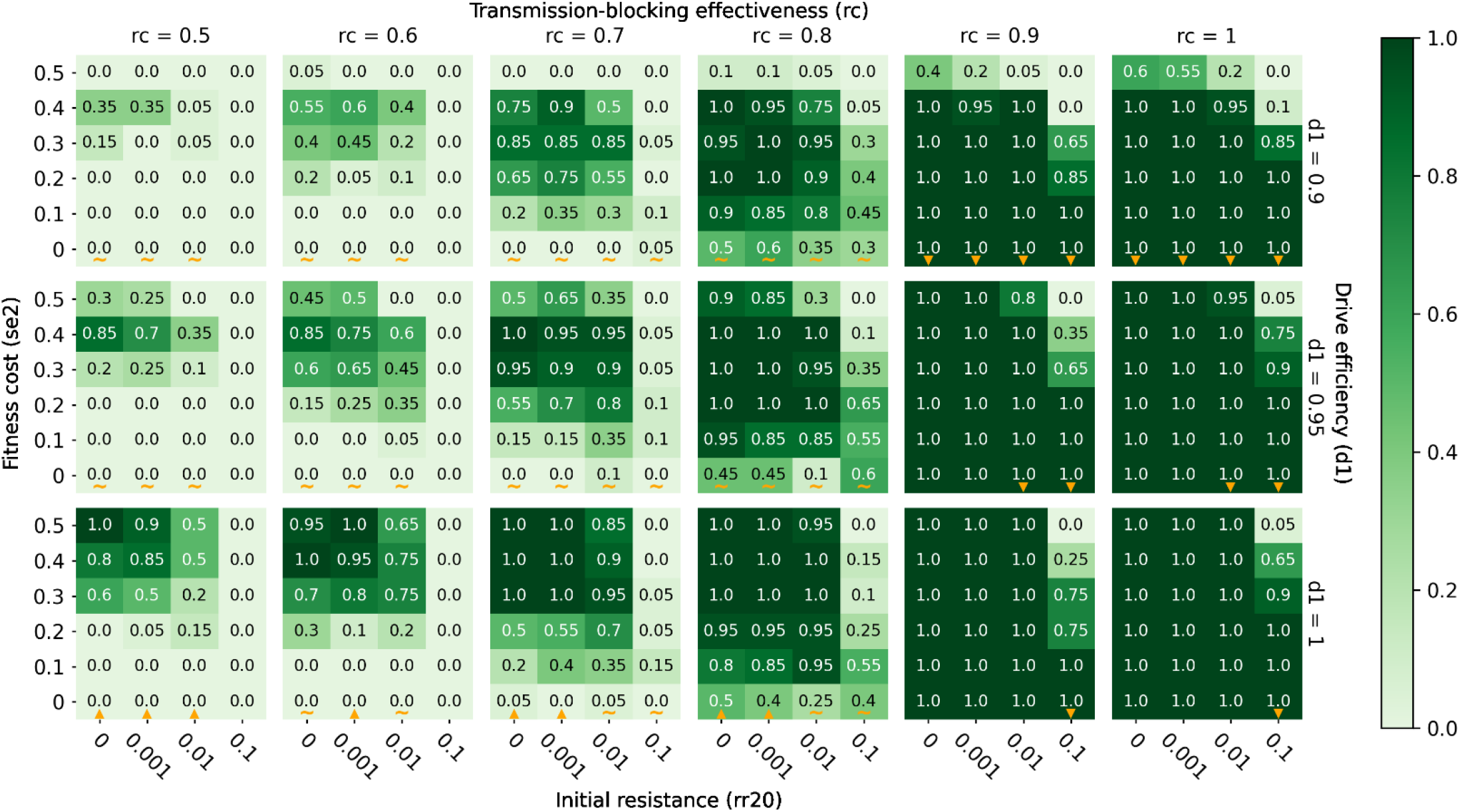
Elimination probabilities after a single release of integral gene drive mosquitoes and ITN deployment in a high transmission (annual EIR = 80) regime. Same as Figure 3.

## Acknowledgments

The authors would like to thank Svetlana Titova, Daniel Bridenbecker, and Clinton Collins for model and software support. We would also like to thank Bee Workeneh, David Kong, Dejan Lukacevic, and Clinton Collins for the tremendous amount of work they did to help us set up the interactive visualization website.

## Author contributions

SL: Conceptualization, Formal analysis, Investigation, Methodology, Project administration, Software, Supervision, Validation, Visualization, Writing – original draft, Writing – review & editing

NW: Writing – Review & Editing

EW: Writing – Review & Editing

CB: Writing – Review & Editing

PS: Conceptualization, Methodology, Project administration, Software, Supervision, Visualization, Writing – original draft, Writing – review & editing

## Data availability

The input files, model executable, and code for running simulations as well as analyzing and plotting model output can be found on Github (https://github.com/InstituteforDiseaseModeling/leung-gene-drive-2021). Software dependencies such as dtk-tools, dtk-tools-malaria, and the malaria-toolbox packages are available upon request from.

## References

1. World malaria report 2020 - 20 years of global progress & challenges. : 300.

2. Bhatt S, Weiss DJ, Cameron E, Bisanzio D, Mappin B, Dalrymple U, et al. The effect of malaria control on Plasmodium falciparum in Africa between 2000 and 2015. Nature. 2015;526: 207–211. doi:10.1038/nature15535

3. Feachem RGA, Chen I, Akbari O, Bertozzi-Villa A, Bhatt S, Binka F, et al. Malaria eradication within a generation: ambitious, achievable, and necessary. The Lancet. 2019;394: 1056–1112. doi:10.1016/S0140-6736(19)31139-0

4. Cook J, Tomlinson S, Kleinschmidt I, Donnelly MJ, Akogbeto M, Adechoubou A, et al. Implications of insecticide resistance for malaria vector control with long-lasting insecticidal nets: trends in pyrethroid resistance during a WHO-coordinated multi-country prospective study. Parasites Vectors. 2018;11: 550. doi:10.1186/s13071-018-3101-4

5. Hancock PA, Hendriks CJM, Tangena J-A, Gibson H, Hemingway J, Coleman M, et al. Mapping trends in insecticide resistance phenotypes in African malaria vectors. PLOS Biology. 2020;18: e3000633. doi:10.1371/journal.pbio.3000633

6. Hemingway J, Ranson H, Magill A, Kolaczinski J, Fornadel C, Gimnig J, et al. Averting a malaria disaster: will insecticide resistance derail malaria control? The Lancet. 2016;387: 1785–1788. doi:10.1016/S0140-6736(15)00417-1

7. Sougoufara S, Doucoure S, Sembéne P, Harry M, Sokhna C. Challenges for malaria vector control in sub-Saharan Africa: Resistance and behavioral adaptations in Anopheles populations. Journal of Vector Borne Diseases. 2017;54: 4–15.

8. Carvalho DO, McKemey AR, Garziera L, Lacroix R, Donnelly CA, Alphey L, et al. Suppression of a Field Population of Aedes aegypti in Brazil by Sustained Release of Transgenic Male Mosquitoes. PLOS Neglected Tropical Diseases. 2015;9: e0003864. doi:10.1371/journal.pntd.0003864

9. Alphey L, McKemey A, Nimmo D, Neira Oviedo M, Lacroix R, Matzen K, et al. Genetic control of Aedes mosquitoes. Pathogens and Global Health. 2013;107: 170–179. doi:10.1179/2047773213Y.0000000095

10. Harris AF, McKemey AR, Nimmo D, Curtis Z, Black I, Morgan SA, et al. Successful suppression of a field mosquito population by sustained release of engineered male mosquitoes. Nat Biotechnol. 2012;30: 828–830. doi:10.1038/nbt.2350

11. Lacroix R, McKemey AR, Raduan N, Wee LK, Ming WH, Ney TG, et al. Open Field Release of Genetically Engineered Sterile Male Aedes aegypti in Malaysia. PLOS ONE. 2012;7: e42771. doi:10.1371/journal.pone.0042771

12. Harris AF, Nimmo D, McKemey AR, Kelly N, Scaife S, Donnelly CA, et al. Field performance of engineered male mosquitoes. Nat Biotechnol. 2011;29: 1034–1037. doi:10.1038/nbt.2019

13. Adolfi A, Gantz VM, Jasinskiene N, Lee H-F, Hwang K, Terradas G, et al. Efficient population modification gene-drive rescue system in the malaria mosquito Anopheles stephensi. Nat Commun. 2020;11: 5553. doi:10.1038/s41467-020-19426-0

14. Carballar-Lejarazú R, James AA. Population modification of Anopheline species to control malaria transmission. Pathogens and Global Health. 2017;111: 424–435. doi:10.1080/20477724.2018.1427192

15. Champer J, Buchman A, Akbari OS. Cheating evolution: engineering gene drives to manipulate the fate of wild populations. Nat Rev Genet. 2016;17: 146–159. doi:10.1038/nrg.2015.34

16. Burt A. Site-specific selfish genes as tools for the control and genetic engineering of natural populations. Proceedings of the Royal Society of London Series B: Biological Sciences. 2003 [cited 16 Aug 2021]. doi:10.1098/rspb.2002.2319

17. James A, Beerntsen B, Capurro M, Coates C, Coleman J, Jasinskiene N, et al. Controlling malaria transmission with genetically-engineered, Plasmodium-resistant mosquitoes: Milestones in a model system. Parassitologia. 1999;41: 461–71.

18. Wang S, Jacobs-Lorena M. Genetic approaches to interfere with malaria transmission by vector mosquitoes. Trends in Biotechnology. 2013;31: 185–193. doi:10.1016/j.tibtech.2013.01.001

19. Isaacs AT, Li F, Jasinskiene N, Chen X, Nirmala X, Marinotti O, et al. Engineered Resistance to Plasmodium falciparum Development in Transgenic Anopheles stephensi. PLOS Pathogens. 2011;7: e1002017. doi:10.1371/journal.ppat.1002017

20. Isaacs AT, Jasinskiene N, Tretiakov M, Thiery I, Zettor A, Bourgouin C, et al. Transgenic Anopheles stephensi coexpressing single-chain antibodies resist Plasmodium falciparum development. PNAS. 2012;109: E1922–E1930. doi:10.1073/pnas.1207738109

21. Ito J, Ghosh A, Moreira LA, Wimmer EA, Jacobs-Lorena M. Transgenic anopheline mosquitoes impaired in transmission of a malaria parasite. Nature. 2002;417: 452–455. doi:10.1038/417452a

22. Corby-Harris V, Drexler A, Jong LW de, Antonova Y, Pakpour N, Ziegler R, et al. Activation of Akt Signaling Reduces the Prevalence and Intensity of Malaria Parasite Infection and Lifespan in Anopheles stephensi Mosquitoes. PLOS Pathogens. 2010;6: e1001003. doi:10.1371/journal.ppat.1001003

23. Adelman ZN, Kojin BB. Malaria-Resistant Mosquitoes (Diptera: Culicidae); The Principle is Proven, But Will the Effectors Be Effective? Journal of Medical Entomology. 2021 [cited 28 Aug 2021]. doi:10.1093/jme/tjab090

24. Champer J, Reeves R, Oh SY, Liu C, Liu J, Clark AG, et al. Novel CRISPR/Cas9 gene drive constructs reveal insights into mechanisms of resistance allele formation and drive efficiency in genetically diverse populations. PLOS Genetics. 2017;13: e1006796. doi:10.1371/journal.pgen.1006796

25. Hammond AM, Kyrou K, Bruttini M, North A, Galizi R, Karlsson X, et al. The creation and selection of mutations resistant to a gene drive over multiple generations in the malaria mosquito. PLOS Genetics. 2017;13: e1007039. doi:10.1371/journal.pgen.1007039

26. Champer J, Liu J, Oh SY, Reeves R, Luthra A, Oakes N, et al. Reducing resistance allele formation in CRISPR gene drive. PNAS. 2018; 115: 5522–5527. doi:10.1073/pnas.1720354115

27. Kranjc N, Crisanti A, Nolan T, Bernardini F. Anopheles gambiae Genome Conservation as a Resource for Rational Gene Drive Target Site Selection. Insects. 2021;12: 97. doi:10.3390/insects12020097

28. James S, Collins FH, Welkhoff PA, Emerson C, Godfray HCJ, Gottlieb M, et al. Pathway to Deployment of Gene Drive Mosquitoes as a Potential Biocontrol Tool for Elimination of Malaria in Sub-Saharan Africa: Recommendations of a Scientific Working Group. The American Journal of Tropical Medicine and Hygiene. 2018;98: 1–49. doi:10.4269/ajtmh.18-0083

29. Brossard D, Belluck P, Gould F, Wirz CD. Promises and perils of gene drives: Navigating the communication of complex, post-normal science. PNAS. 2019;116: 7692–7697. doi:10.1073/pnas.1805874115

30. Connolly JB, Mumford JD, Fuchs S, Turner G, Beech C, North AR, et al. Systematic identification of plausible pathways to potential harm via problem formulation for investigational releases of a population suppression gene drive to control the human malaria vector Anopheles gambiae in West Africa. Malaria Journal. 2021;20: 170. doi:10.1186/s12936-021-03674-6

31. Beaghton A, Hammond A, Nolan T, Crisanti A, Godfray HCJ, Burt A. Requirements for Driving Antipathogen Effector Genes into Populations of Disease Vectors by Homing. Genetics. 2017;205: 1587–1596. doi:10.1534/genetics.116.197632

32. Selvaraj P, Wenger EA, Bridenbecker D, Windbichler N, Russell JR, Gerardin J, et al. Vector genetics, insecticide resistance and gene drives: An agent-based modeling approach to evaluate malaria transmission and elimination. Davenport MP, editor. PLoS Comput Biol. 2020;16: e1008121. doi:10.1371/journal.pcbi.1008121

33. North AR, Burt A, Godfray HCJ. Modelling the suppression of a malaria vector using a CRISPR-Cas9 gene drive to reduce female fertility. BMC Biology. 2020;18: 98. doi:10.1186/s12915-020-00834-z

34. North A, Burt A, Godfray HCJ. Modelling the spatial spread of a homing endonuclease gene in a mosquito population. Journal of Applied Ecology. 2013;50: 1216–1225. doi:10.1111/1365-2664.12133

35. C HMS, Wu SL, Bennett JB, Marshall JM. MGDrivE: A modular simulation framework for the spread of gene drives through spatially explicit mosquito populations. Methods in Ecology and Evolution. 2020;11: 229–239. doi:10.1111/2041-210X.13318

36. Marshall JM, Buchman A, Sánchez C. HM, Akbari OS. Overcoming evolved resistance to population-suppressing homing-based gene drives. Sci Rep. 2017;7: 3776. doi:10.1038/s41598-017-02744-7

37. Nash A, Urdaneta GM, Beaghton AK, Hoermann A, Papathanos PA, Christophides GK, et al. Integral gene drives for population replacement. Biology Open. 2018; bio.037762. doi:10.1242/bio.037762

38. Eckhoff PA, Wenger EA, Godfray HCJ, Burt A. Impact of mosquito gene drive on malaria elimination in a computational model with explicit spatial and temporal dynamics. Proc Natl Acad Sci USA. 2017;114: E255–E264. doi:10.1073/pnas.1611064114

39. Epidemiological Modeling Software. Institute for Disease Modeling; 2021. Available: http://idmod.org

40. Eckhoff PA. A malaria transmission-directed model of mosquito life cycle and ecology. Malar J. 2011;10: 303. doi:10.1186/1475-2875-10-303

41. Selvaraj P, Wenger EA, Gerardin J. Seasonality and heterogeneity of malaria transmission determine success of interventions in high-endemic settings: a modeling study. BMC Infectious Diseases. 2018;18: 413. doi:10.1186/s12879-018-3319-y

42. Center for International Earth Science Information Network. [cited 16 Aug 2021]. Available: https://www.ciesin.columbia.edu/data/hrsl/

43. Molineaux L, Gramiccia G. The Garki Project. Research on the epidemiology and control of malaria in the Sudan savanna of West Africa. Transactions of the Royal Society of Tropical Medicine and Hygiene. 1981;75: 190–191. doi:10.1016/0035-9203(81)90085-7

44. Huestis DL, Dao A, Diallo M, Sanogo ZL, Samake D, Yaro AS, et al. Windborne long-distance migration of malaria mosquitoes in the Sahel. Nature. 2019;574: 404–408. doi:10.1038/s41586-019-1622-4

45. Collins KA, Ouedraogo A, Guelbeogo WM, Awandu SS, Stone W, Soulama I, et al. Investigating the impact of enhanced community case management and monthly screening and treatment on the transmissibility of malaria infections in Burkina Faso: study protocol for a cluster-randomised trial. BMJ Open. 2019;9: e030598. doi:10.1136/bmjopen-2019-030598

46. Selvaraj P, Suresh J, Wenger EA, Bever CA, Gerardin J. Reducing malaria burden and accelerating elimination with long-lasting systemic insecticides: a modelling study of three potential use cases. Malaria Journal. 2019;18: 307. doi:10.1186/s12936-019-2942-4

47. Eisele TP, Bennett A, Silumbe K, Finn TP, Chalwe V, Kamuliwo M, et al. Short-term Impact of Mass Drug Administration With Dihydroartemisinin Plus Piperaquine on Malaria in Southern Province Zambia: A Cluster-Randomized Controlled Trial. The Journal of Infectious Diseases. 2016;214: 1831–1839. doi:10.1093/infdis/jiw416

48. Thomas CJ, Cross DE, Bøgh C. Landscape Movements of Anopheles gambiae Malaria Vector Mosquitoes in Rural Gambia. Shiff C, editor. PLoS ONE. 2013;8: e68679. doi:10.1371/journal.pone.0068679

49. Achieving and maintaining universal coverage with long-lasting insecticidal nets for malaria control. : 4.

50. Schmidt H, Collier TC, Hanemaaijer MJ, Houston PD, Lee Y, Lanzaro GC. Abundance of conserved CRISPR-Cas9 target sites within the highly polymorphic genomes of Anopheles and Aedes mosquitoes. Nat Commun. 2020;11: 1425. doi:10.1038/s41467-020-15204-0

51. Carballar-Lejarazú R, Ogaugwu C, Tushar T, Kelsey A, Pham TB, Murphy J, et al. Next-generation gene drive for population modification of the malaria vector mosquito, Anopheles gambiae. PNAS. 2020;117: 22805–22814. doi:10.1073/pnas.2010214117

52. Kyrou K, Hammond AM, Galizi R, Kranjc N, Burt A, Beaghton AK, et al. A CRISPR–Cas9 gene drive targeting doublesex causes complete population suppression in caged Anopheles gambiae mosquitoes. Nat Biotechnol. 2018;36: 1062–1066. doi:10.1038/nbt.4245

53. Gantz VM, Jasinskiene N, Tatarenkova O, Fazekas A, Macias VM, Bier E, et al. Highly efficient Cas9-mediated gene drive for population modification of the malaria vector mosquito Anopheles stephensi. PNAS. 2015;112: E6736–E6743. doi:10.1073/pnas.1521077112

54. Hammond A, Galizi R, Kyrou K, Simoni A, Siniscalchi C, Katsanos D, et al. A CRISPR-Cas9 gene drive system targeting female reproduction in the malaria mosquito vector Anopheles gambiae. Nat Biotechnol. 2016;34: 78–83. doi:10.1038/nbt.3439

55. Pham TB, Phong CH, Bennett JB, Hwang K, Jasinskiene N, Parker K, et al. Experimental population modification of the malaria vector mosquito, Anopheles stephensi. PLOS Genetics. 2019;15: e1008440. doi:10.1371/journal.pgen.1008440

56. McArthur CC, Meredith JM, Eggleston P. Transgenic Anopheles gambiae Expressing an Antimalarial Peptide Suffer No Significant Fitness Cost. PLOS ONE. 2014;9: e88625. doi:10.1371/journal.pone.0088625

57. Dong S, Fu X, Dong Y, Simões ML, Zhu J, Dimopoulos G. Broad spectrum immunomodulatory effects of Anopheles gambiae microRNAs and their use for transgenic suppression of Plasmodium. PLOS Pathogens. 2020;16: e1008453. doi:10.1371/journal.ppat.1008453

58. Hoermann A, Tapanelli S, Capriotti P, Del Corsano G, Masters EK, Habtewold T, et al. Converting endogenous genes of the malaria mosquito into simple non-autonomous gene drives for population replacement. Messer PW, Tautz D, Lawniczak M, Marois E, editors. eLife. 2021;10: e58791. doi:10.7554/eLife.58791

59. Dong Y, Simões ML, Dimopoulos G. Versatile transgenic multistage effector-gene combinations for Plasmodium falciparum suppression in Anopheles. Science Advances. 2020 [cited 1 Sep 2021]. Available: https://www.science.org/doi/abs/10.1126/sciadv.aay5898

60. Dong Y, Das S, Cirimotich C, Souza-Neto JA, McLean KJ, Dimopoulos G. Engineered Anopheles Immunity to Plasmodium Infection. PLOS Pathogens. 2011;7: e1002458. doi:10.1371/journal.ppat.1002458

61. Meredith JM, Basu S, Nimmo DD, Larget-Thiery I, Warr EL, Underhill A, et al. Site-Specific Integration and Expression of an Anti-Malarial Gene in Transgenic Anopheles gambiae Significantly Reduces Plasmodium Infections. PLOS ONE. 2011;6: e14587. doi:10.1371/journal.pone.0014587

62. Volohonsky G, Hopp A-K, Saenger M, Soichot J, Scholze H, Boch J, et al. Transgenic Expression of the Anti-parasitic Factor TEP1 in the Malaria Mosquito Anopheles gambiae. PLOS Pathogens. 2017;13: e1006113. doi:10.1371/journal.ppat.1006113

63. James SL, Marshall JM, Christophides GK, Okumu FO, Nolan T. Toward the Definition of Efficacy and Safety Criteria for Advancing Gene Drive-Modified Mosquitoes to Field Testing. Vector-Borne and Zoonotic Diseases. 2020;20: 237–251. doi:10.1089/vbz.2019.2606

